# Transcriptome-wide alternative mRNA splicing analysis reveals post-transcriptional regulation of neuronal differentiation

**DOI:** 10.1101/2024.07.16.603656

**Authors:** Yuan Zhou, Sherif Rashad, Kuniyasu Niizuma

## Abstract

Alternative splicing (AS) plays important roles in neuronal development, function, and diseases. Efforts to analyze AS transcriptome-wide in neurons remain limited. We characterized the transcriptome-wide AS changes in SH-SY5Y neuronal differentiation model, which is widely used to study neuronal function and disorders. Our analysis revealed global changes in five AS programs that drive neuronal differentiation. Motif analysis revealed the contribution of RNA binding proteins (RBPs) to the regulation of AS during neuronal development. We focused on the predominant AS program during differentiation, exon skipping (SE) events. Motif analysis revealed motifs for PTB and HuR/ELAVL1 to be the top enriched in SE events, and their protein levels were downregulated after differentiation. shRNA Knockdown of either PTB and HuR were associated with enhanced neuronal differentiation and transcriptome-wide exon skipping events driving the process of differentiation. At the level of gene expression, we observed only modest changes, indicating predominant post-transcriptional effects of PTB and HuR. We also observed that both RBPs altered cellular responses to oxidative stress, in line with the differentiated phenotype observed after KD. Our work characterizes the AS changes in a widely used and important model of neuronal development and neuroscience research and reveals intricate post-transcriptional regulation of neuronal differentiation.

## Introduction

Neuronal differentiation is a complex multistep process in which neurons undergo dramatic morphological alterations, including neurite outgrowth and synapse formation(1). A series of changes then enables neuron to carry out their activities such as, electrophysiological activity and neurotransmitter secretion (2,3). Consequently, these changes render mature neurons highly sensitive to oxidative stress, a property which plays a crucial etiology in many neurological and neurodegenerative diseases (4–6). Given the important link between oxidative stress in neurons and neurodegenerative diseases such as Alzheimer’s(7) and Parkinson’s(8), it is crucial not only in understanding the pathophysiology of diseases, but also to develop strategies for neuroprotection via genetic or pharmacological intervention.

Recent advances have revealed an important role that the epigenome plays in regulating neuronal behavior as well as neurogenesis. For example, a dynamic mRNA decay network regulated by RNA binding proteins (RBPs) Pumilio contributes to the relative abundance of transcripts involved in cell-fate decisions and axonogenesis during Drosophila neural development(9). Moreover, emerging evidence unveiled a strong association between altered mRNA stability and neurodegenerative disease (10). On the other hand, neurons at their different developmental stages have specific alternative splicing (AS) patterns, indicating a potential association of AS in regulating neuronal differentiation and maintaining neuron maturation(11). While many post-transcriptional events occurring during neuronal differentiation are now increasingly understood, it remains largely unknown how exactly post-transcriptional regulation contributes to neuronal differentiation. Furthermore, how the changes associated with neuronal differentiation and maturation render neurons more sensitive to oxidative stress is not fully understood.

SH-SY5Y human neuroblastoma cells are frequently used as *in vitro* neuronal differentiation model due to the biochemical and morphological resemblance of neuron to study neuronal physiology and neurodegenerative diseases(2,12). Previous transcriptomic profiling on both differentiated and undifferentiated SH-SY5Y cells revealed that differential gene expression in these neuron-like cells are more functionally linked to changes in cell morphology including remodeling of plasma membrane and cytoskeleton, and neuritogenesis (6,13,14). On the other hand, SH-SY5Y cells develop distinct responses to cellular stressors/neurotoxins after differentiation which leads to their utilization as cell model systems to study the mechanisms of neurodegenerative disorders (15). However, existing transcriptomic analysis on differentiated SH-SY5Y cells cannot provide molecular basis to this heightened sensitivity to stress(13,14), nor can it account for the entirety of the neurogenic process(6).

Previously, we showed that during SH-SY5Y differentiation, there are transcriptome-wide changes in mRNA stability that are driven by RBPs (6). SAMD4A was identified as one of the regulators of this process and in turn, of neuronal differentiation and stress response to oxidative and mitochondrial stress. In this work, we focused on another aspect of post-transcriptional changes that occur during neuronal differentiation, alternative splicing. We aimed to characterize the transcriptome-wide changes occurring in mRNA alternative splicing during neuronal differentiation, and to identify potential regulators. We show that during neuronal differentiation there are significant changes in multiple AS programs that drive neuronal differentiation. Interrupting these programs by targeting the regulatory RBPs impacts neuronal differentiation. These results add an additional layer of understanding to the process of neuronal differentiation, and the implications in understanding neural plasticity, development, and neurological disorders.

## Results

### Transcriptome-wide analysis of local splice variates (LSVs) during neuronal differentiation

We previously conducted RNA-seq analysis on differentiated and undifferentiated SH-SY5Y cells (6). In this previous work, the differentiation of SH-SY5Y cells was validated by multiple methods prior to sequencing. To analyze alternative splicing, we employed the rMATs pipeline (16) to analyze local splice variants (LSV). We evaluated mRNA LSVs in 5 different splicing programs; skipped exon (SE), mutually exclusive exons (MXE), retained introns (RI), alternative 5’ splice site (A5SS), and alternative 3’ splice site (A3SS). Our analysis revealed thousands of AS events in 2557 unique genes in each of these programs, with SE representing the largest number of AS events and alternatively spliced genes [Figure 1A-G, Supplementary tables 1-5]. Next, we selected the significantly spliced genes from each program (FDR < 0.05, ΔΨ >0.2 or <-0.2) and conducted pathway overrepresentation analysis (ORA) [Figure 1H]. Gene ontology biological processes (GOBP) analysis revealed significant enrichment of pathways involved in cellular replication, morphogenesis, and cytoskeletal organization, in agreement with the changes expected during neuronal differentiation as well as with the changes observed at the level of differential gene expression (6). SE was the most enriched in these pathways, however, we observed enrichment of translation linked pathways that were more enriched in other LSV programs such as RI or A3SS.

**Figure 1:**
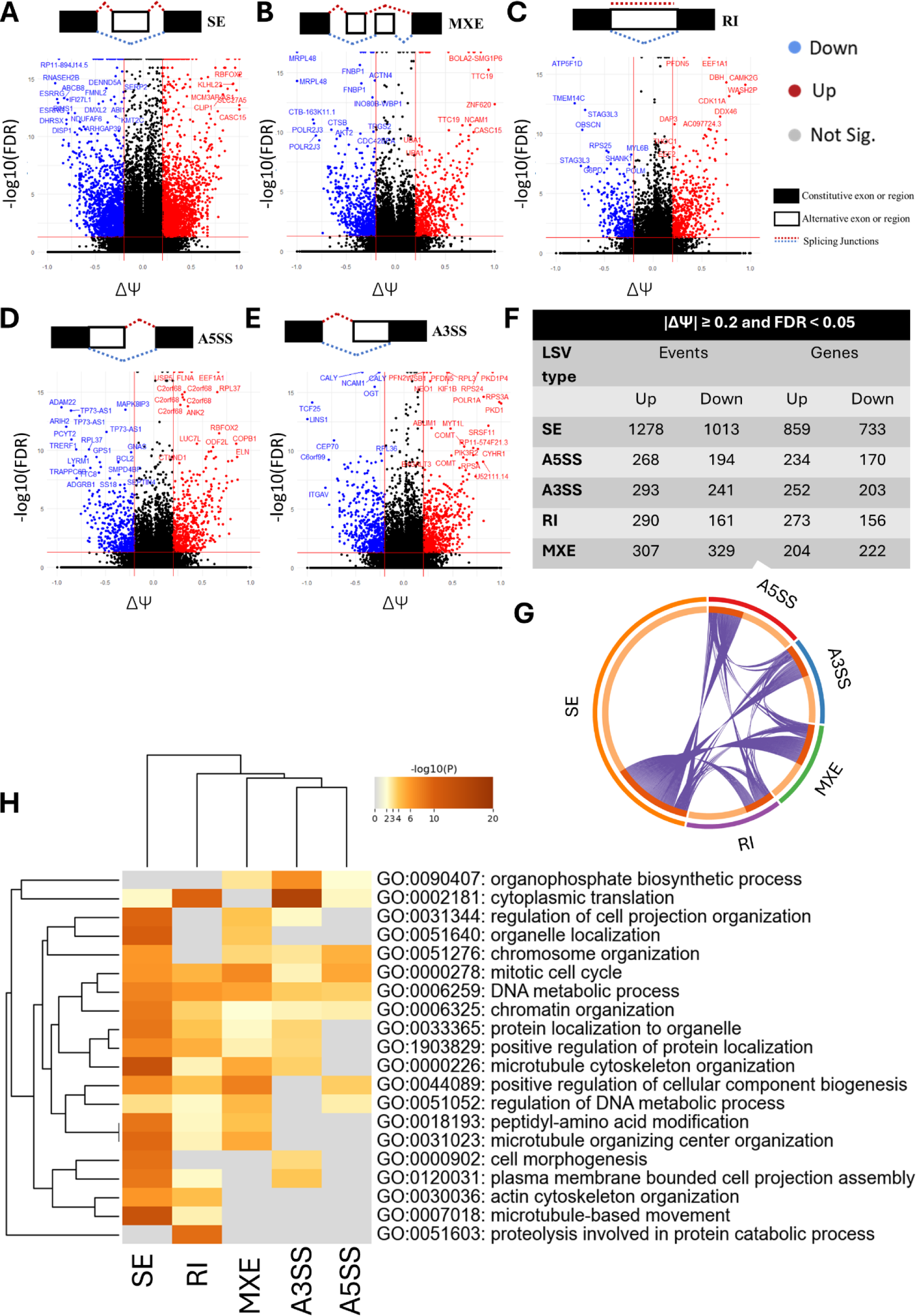
Transcriptome-wide analysis of alternative splicing changes during neuronal differentiation. **A-E:** Volcano plots representing the various local splice variant (LSV) programs activated during neuronal differentiation. **F:** Summary of the number of significant events and alternatively spliced genes in each LSV program. **G:** Circos plot showing the overlap in genes detected in each LSV program. **H:** Overrepresentation analysis (ORA) showing the enrichment in GOBP terms in each LSV program.

### LSVs are regulated by RNA binding proteins

AS is regulated by many factors, including the spliceosome components, mRNA modifications, and RNA binding proteins (RBPs) acting as splice regulators (17–19). To that end, we explored our previous RNA-seq and Ribo-seq for the expression of spliceosome genes and RBPs related to mRNA alternative splicing (the list of RBPs was compiled from (20,21) and retrieved from rMAPs2 database (22)). First, we observed global downregulation of the spliceosome pathway at the level of RNA expression [Supplementary figure 1A-B] and RNA translation [Supplementary figure 1C-D]. Further, analysis of RBPs gene expression [Supplementary figure 1E] and mRNA translation levels [Supplementary figure 1F] revealed changes in several RBPs. Importantly, we observed upregulation of SAMD4A, CELF6, and CPEB2 and downregulation of SNRPA, RBP42, and PABPC3 at the level of mRNA translation. We previously showed that SAMD4A, which was upregulated in the RNA-seq and Ribo-seq datasets, is upregulated during neuronal differentiation at the protein level, that it regulates mRNA stability, and that it regulates neuronal differentiation and response to mitochondrial stress and ferroptosis induction (6).

Given that our analysis pointed to global AS changes, we directed our attention to how RBPs might influence the global changes at the level of LSVs observed. To that end, we conducted motif search on the LSV data using rMAPs2 software (22). Motif enrichment revealed global changes in the motifs of RBPs that could direct multiple LSV programs during neuronal differentiation. For example, PTB/PTBP1 motif was the top enriched in the upregulated SE events after differentiation [Figure 2A and C], while multiple HuR/ELAVL1 motifs were enriched in the downregulated SE events [Figure 2B and D, supplementary figure 2]. In RI events, we observed HNRNPL in the upregulated events to be the top enriched, while ZC3H10 and PTB to be the top enriched in the downregulated events [Supplementary figure 3]. KHDRBS3 and RBM45 were the top enriched motifs in A5SS [Supplementary figure 4], while SAMD4A and KHDRBS1 were the top enriched motifs in the A3SS events [Supplementary figure 5].

**Figure 2:**
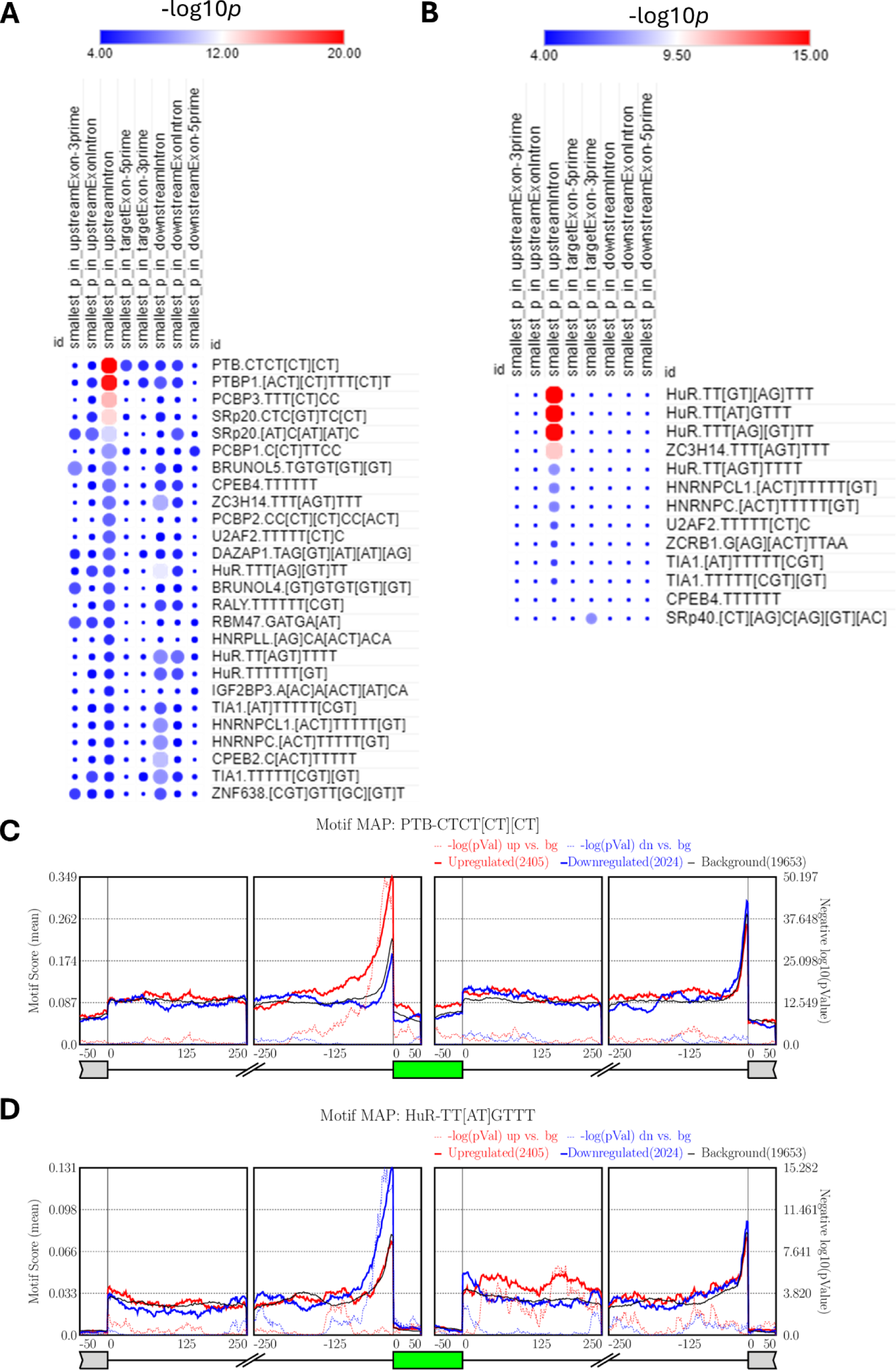
Identification of RBPs regulating exon skipping events using motif analysis. **A:** Heatmap showing the top enriched RBP motifs in the upregulated SE events after neuronal differentiation. **B:** Heatmap showing the top enriched RBP motifs in the downregulated SE events after neuronal differentiation. **C:** Motif-map of PTB motif enrichment. **D:** Motif-map of HuR motif enrichment.

Collectively, our data point to global alterations in AS during neuronal differentiation that could potentially drive the neuronal phenotype and function. Given that we observed a dominance of SE events in our analysis, we opted to validate the results obtained from motif enrichment of SE, selecting the top up and downregulated motifs, PTB and HuR/ELAVL1 respectively for downstream validation.

### PTB regulates neuronal differentiation via modulating mRNA splicing

First, we focused on PTB as a regulator of SE during neuronal differentiation. PTB is known to be repressed during neuronal differentiation and is known to suppress exons skipping (23,24). In line with these observations, we observed that PTB protein levels were downregulated after neuronal differentiation [Figure 3A], in tandem with the enrichment of PTB motif around SE events [Figure 2C]. Next, we designed shRNA to knockdown (KD) PTB in undifferentiation SH-SY5Y cells [Figure 3B]. Upon PTB KD, we observed upregulation of several neuronal differentiation markers such as synuclein (SYN), MAP2, and TUJ1 [Figure 3C]. In addition, when we attempted to differentiate the KD cells, we observed faster, and more robust differentiation and neurite growth compared to the Mock transfected cells [Figure 3D-E]. We also exposed the KD cells to the same stressors that we employed in our previous work, namely, Antimycin A (Respiratory complex III inhibitor), Rotenone (Respiratory complex I inhibitor), Erastin (Class I ferroptosis inducer), RSL3 (Class II ferroptosis inducer), and sodium Arsenite (general oxidative stressor) (6). We also combined our stress analysis with cell death inhibitors to probe which cell death program is activated. We used Ferrostatin-1 (Ferr-1; Ferroptosis inhibitor), Necrostatin-1 (Nec-1; Necroptosis inhibitor), YVAD (Caspase 1/pyroptosis inhibitor), and DEVD (Caspase 3/7 apoptosis inhibitor). As we have previously observed after neuronal differentiation, PTB KD, potentially through driving the cells towards a more differentiated phenotype, sensitized the cells to all stresses used except Arsenite [Figure 3F]. In addition, we observed changes in the programmable cell death (PCD) mechanisms being activated, akin to our observations when we compared differentiated to undifferentiated cells previously (6). For example, In the Mock cells, Erastin activated both Ferroptosis and Pyroptosis, while after PTB KD, Ferroptosis and Necroptosis were activated. Similarly, RSL3 activated Ferroptosis in Mock cells, while after PTB KD it activated Necroptosis and Pyroptosis.

**Figure 3:**
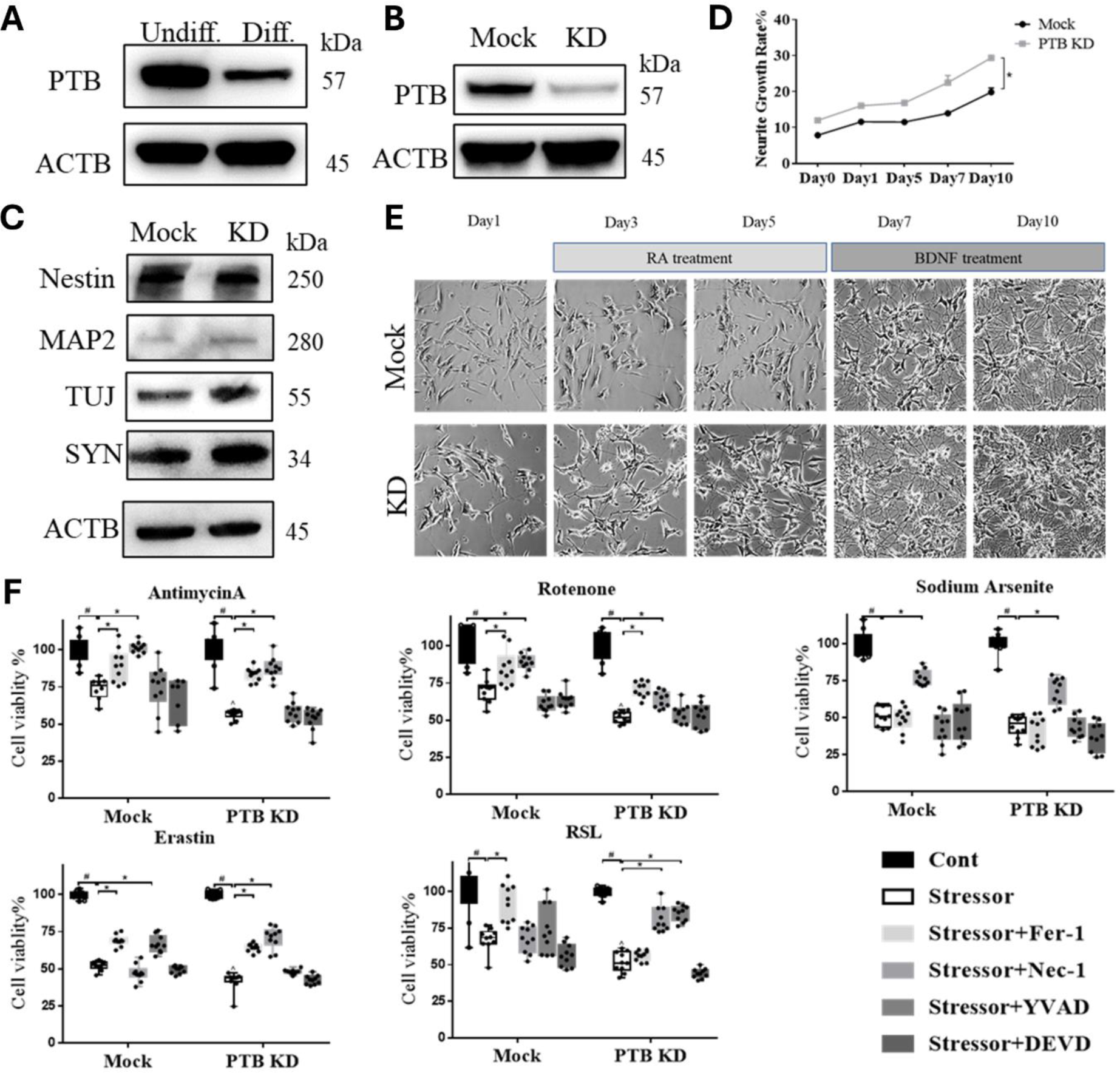
PTB downregulation drives neuronal differentiation. **A:** Western blotting showing the downregulation of PTB after neuronal differentiation. **B:** validation of PTB shRNA knockdown. **C:** Western blotting showing the upregulation of various mature neuronal markers after PTB KD. **D-E:** Phase contrast microscopy analysis of neural differentiation and neurite growth after PTB KD and induction of differentiation. **F:** PTB and Mock KD cells were exposed to 5 different stressors and analyzed using WST-8 assay.

Next, we conducted RNA-seq on the PTB KD cells. Differential gene expression (DEG) revealed modest changes at the level of RNA levels [Figure 4A]. We observed 55 genes to be upregulated and 27 genes to be downregulated after PTB KD (Cutoff criteria: FDR <0.05, Fold change >1.5). Pre-ranked gene set enrichment (GSEA) analysis using GOBP database as reference revealed the upregulation of pathways related to neuronal function such as synaptic activity and neurotransmitter secretion, and the downregulation of pathways related to cell replication [Figure 4B], further validating the shift towards a differentiated phenotype after PTB KD. Next, we analyzed changes in LSVs after PTB KD. We observed hundreds of up and down regulated SE events, as well as other LSV subtypes, after PTB KD [Figure 5A-B, Supplementary tables 6-10]. As expected, SE events were the most dysregulated/altered LSV program. We further analyzed the enrichment of PTB motif after PTB KD. PTB motif was the most upregulated motif in SE events and was significantly enriched around SE events [Figure 5C, supplementary figure 6A]. Analysis of PTB motif enrichment in other LSV subtypes did not reveal significant changes, especially when compared to the strong enrichment in SE [Supplementary figure 6B-E]. Next, we conducted overrepresentation pathway analysis (ORA) on the upregulated SE events, using GOBP as reference, which shows enrichment of pathways related to cytoskeletal organization and neuronal development, such as smoothened signaling (25) and ephrin signaling (26) [Figure 5D]. We also compared the upregulated SE events after PTB with the upregulated SE events after neuronal differentiation, which yielded 166 shared mRNAs [Supplementary figure 7A]. Arguing that those shared mRNAs represent the true effect of PTB during neuronal differentiation amid all the complex changes occurring, we conducted ORA analysis on these 166 genes. ORA analysis revealed the enrichment of many pathways known to play roles in neuronal differentiation and development such smoothened signaling, ephrin signaling, calcium homeostasis, and others [Supplementary figure 7B]. We did not observe any shared genes between the upregulated SE events and the DEGs after PTB KD (not shown).

**Figure 4:**
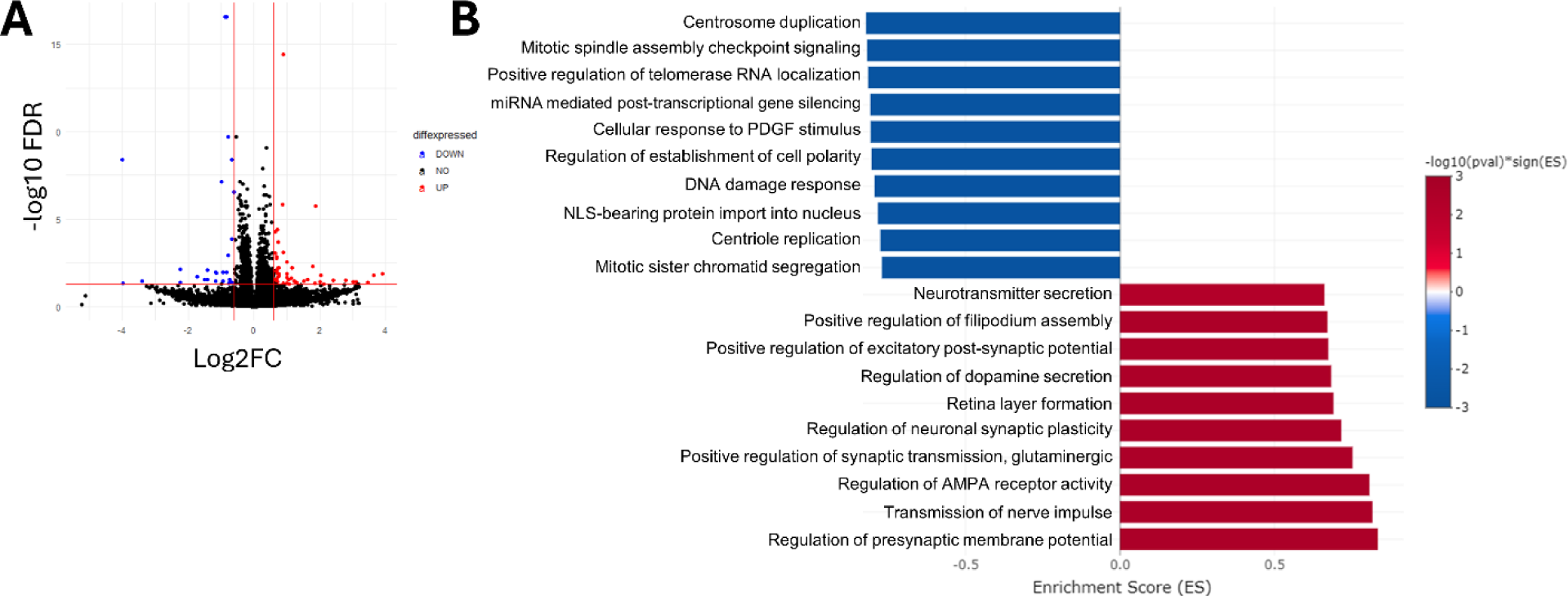
RNA-seq analysis after PTB KD. **A:** Volcano plot showing the differentially expressed genes after PTB KD. **B:** Pre-ranked GSEA showing the enrichment of GOBP terms after PTB KD.

**Figure 5:**
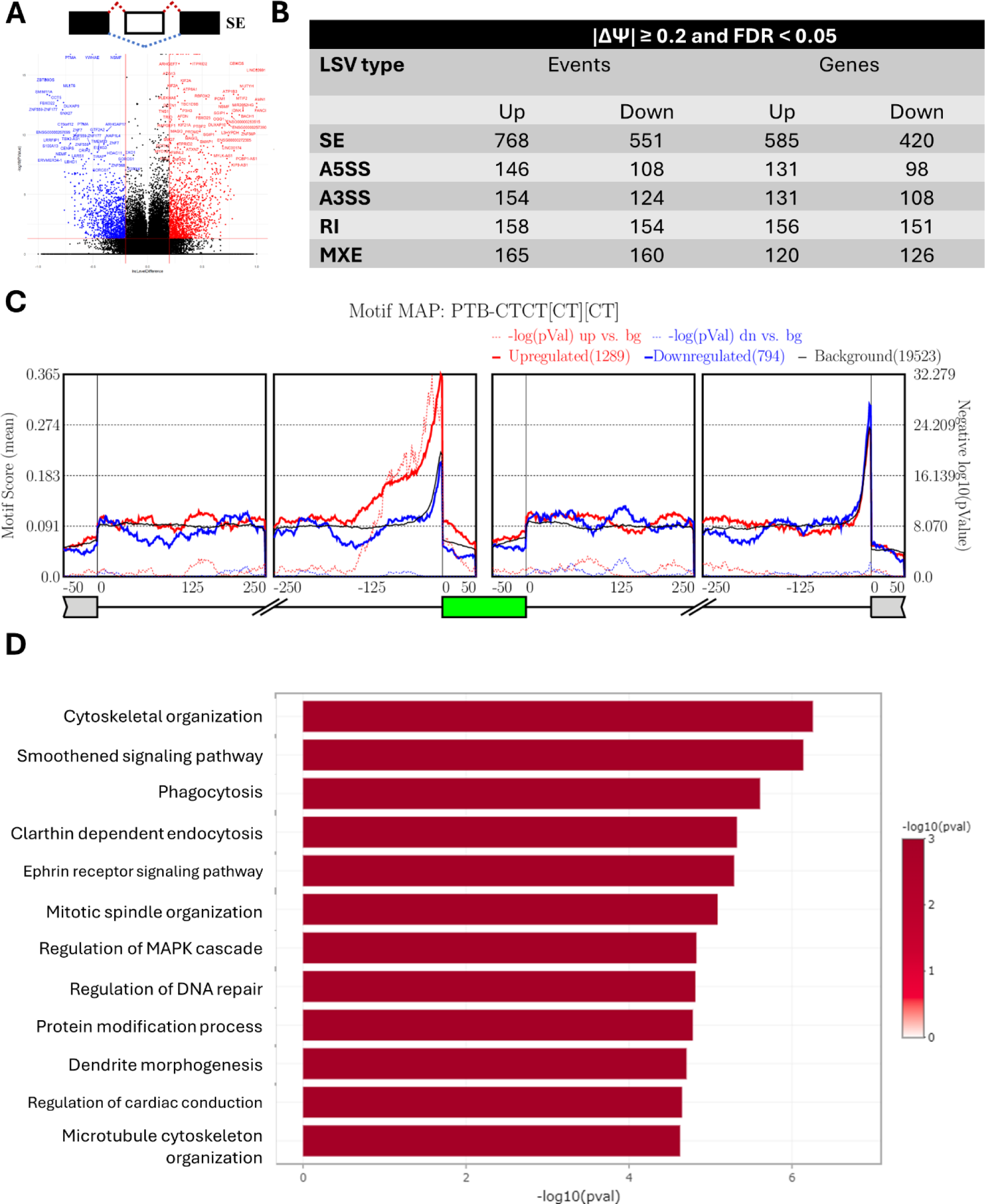
PTB KD induces transcriptome-wide alternative splicing changes. **A:** Volcano plot of skipped exon events after PTB KD. **B:** Table showing the number of significant events and differentially spliced genes in each LSV program after PTB KD. **C:** Motif-map showing the enrichment of PTB motif in the upregulated SE events after PTB KD. **D:** ORA analysis showing the enrichment of GOBP terms in the upregulated SE genes after PTB KD.

In summary, our analysis revealed that PTB downregulation is a driver of neuronal differentiation mainly via promoting exon skipping.

### HuR regulates neuronal differentiation via modulating mRNA splicing

Here, we repeated the same workflow we did above to validate HuR/ELAVL1 role in neuronal differentiation. HuR is known to regulate neuronal development and promote exon skipping (27–29). After neuronal differentiation, HuR protein levels were moderately downregulated [Figure 6A], which agrees with the direction of its motif enrichment observed after neuronal differentiation, where HuR motifs were the top enriched in the downregulated SE events. Next, we knocked down HuR using shRNA [Figure 6B]. HuR KD led to upregulation of the mature neuronal marker SYN [Figure 6C] and led to more robust differentiation of SH-SY5Y cells and faster neurite growth [Figure 6D-E]. Moreover, exposure of HuR KD sensitized cells to stress induction in all stresses employed, except Arsenite, and altered the PCD programs activated in response to stress [Figure 6F].

**Figure 6:**
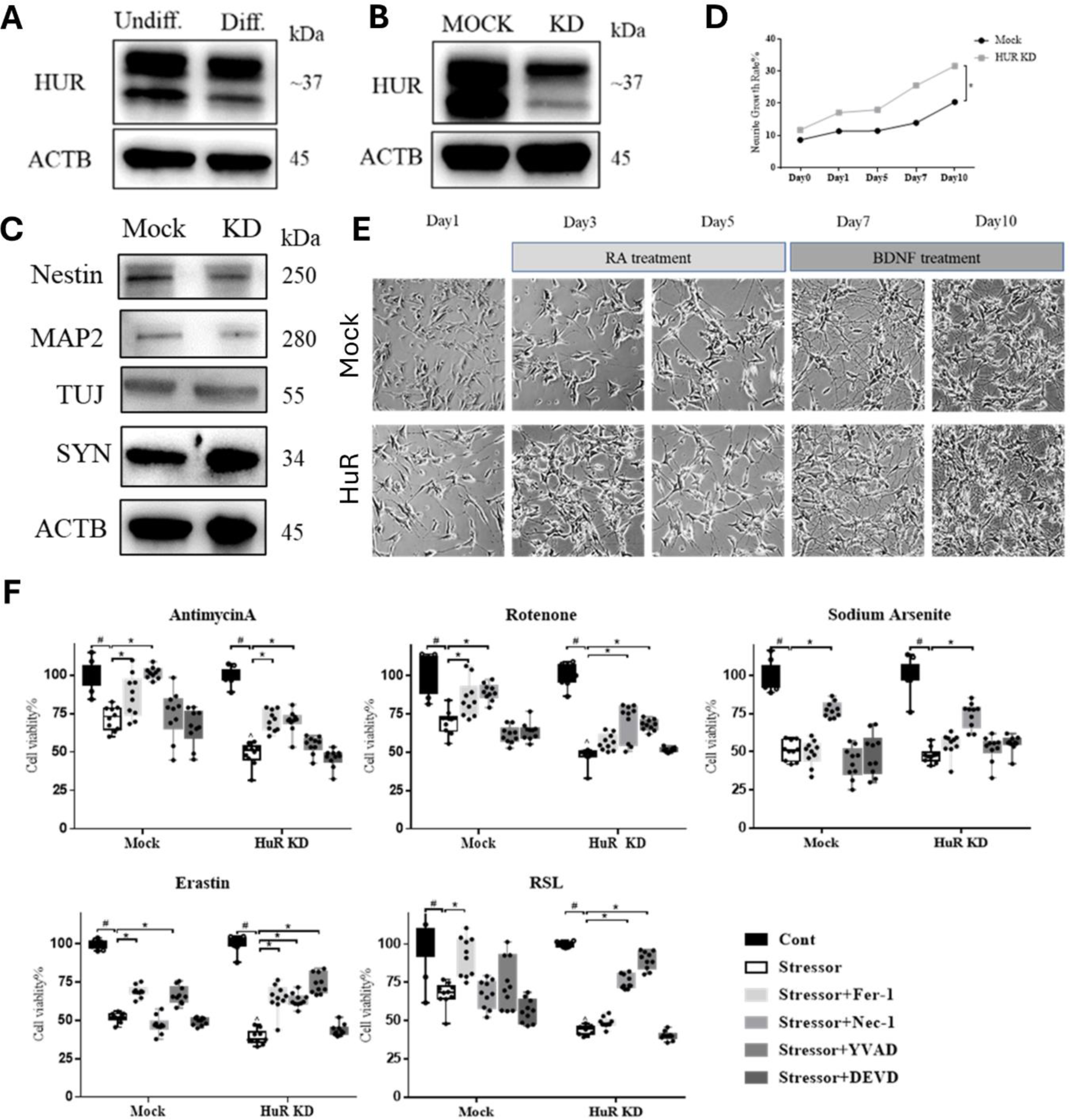
HuR downregulation drives neuronal differentiation. **A:** Western blotting showing the downregulation of HuR after neuronal differentiation. **B:** validation of HuR shRNA knockdown. **C:** Western blotting showing the upregulation of various mature neuronal markers after HuR KD. **D-E:** Phase contrast microscopy analysis of neural differentiation and neurite growth after HuR KD and induction of differentiation. **F:** HuR and Mock KD cells were exposed to 5 different stressors and analyzed using WST-8 assay.

RNA-seq DEG analysis after HuR KD did not show any statistically significant gene expression changes [Figure 7A]. Pre-ranked GSEA analysis revealed the enrichment of pathways related to neuronal differentiation and function and the downregulation of pathways related to cell replication and proliferation [Figure 7B]. On the other hand, we observed a robust response at the level of AS after HuR KD, with SE having the largest number of events/genes [Figure 8A-B, supplementary tables 11-15]. Further, analysis of the multiple HuR motifs revealed enrichment in downregulated SE events, in agreement with HuR’s role in promoting exon skipping [Figure 8C, supplementary figure 8]. Analysis of other LSV motifs did not reveal HuR motif enrichment except in MXE [Supplementary figures 9-12]. We next performed ORA analysis of downregulated SE events after HuR KD [Figure 8D]. GOBP analysis revealed enrichment of pathways related to neuronal function, cellular motility, and protein regulation. Comparing downregulated SE between HuR KD and differentiated cells revealed an overlap of 61 genes [Supplementary figure 13A]. ORA analysis of these 61 genes revealed the enrichment of multiple pathways related to protein activating, neuronal function, and cell cycle [Supplementary figure 13].

**Figure 7:**
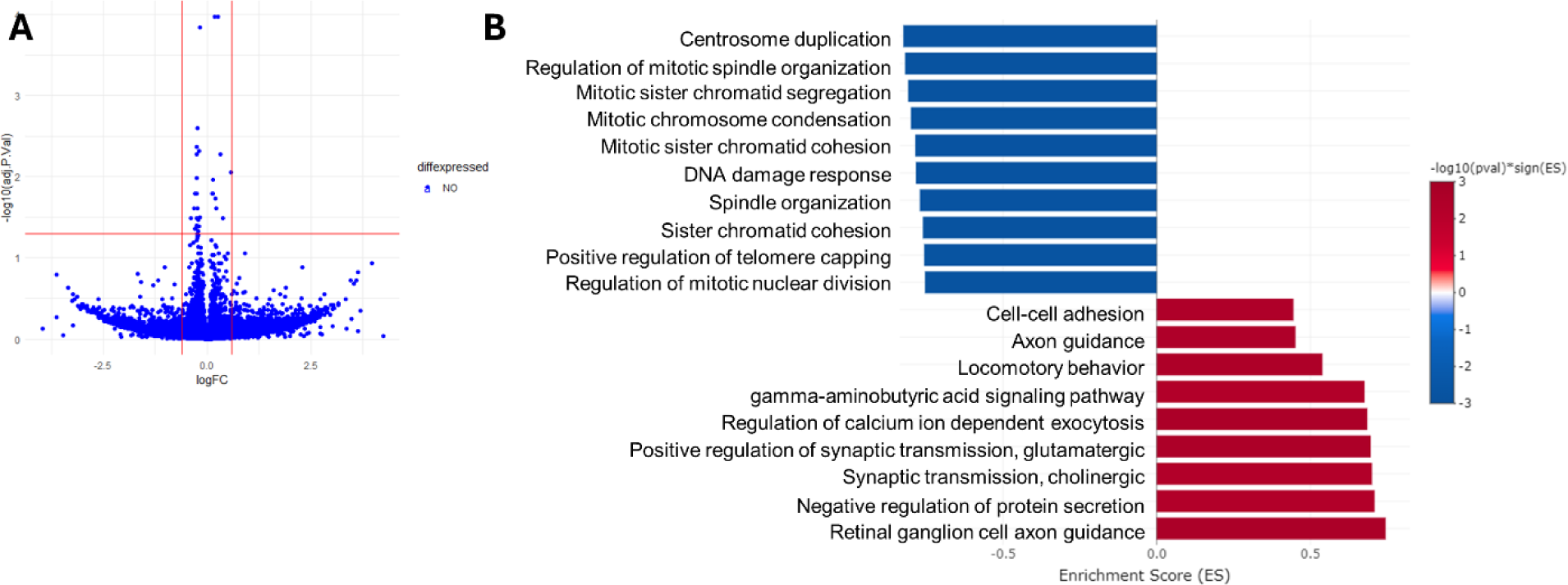
RNA-seq analysis after HuR KD. **A:** Volcano plot showing the differentially expressed genes after HuR KD. **B:** Pre-ranked GSEA showing the enrichment of GOBP terms after HuR KD.

**Figure 8:**
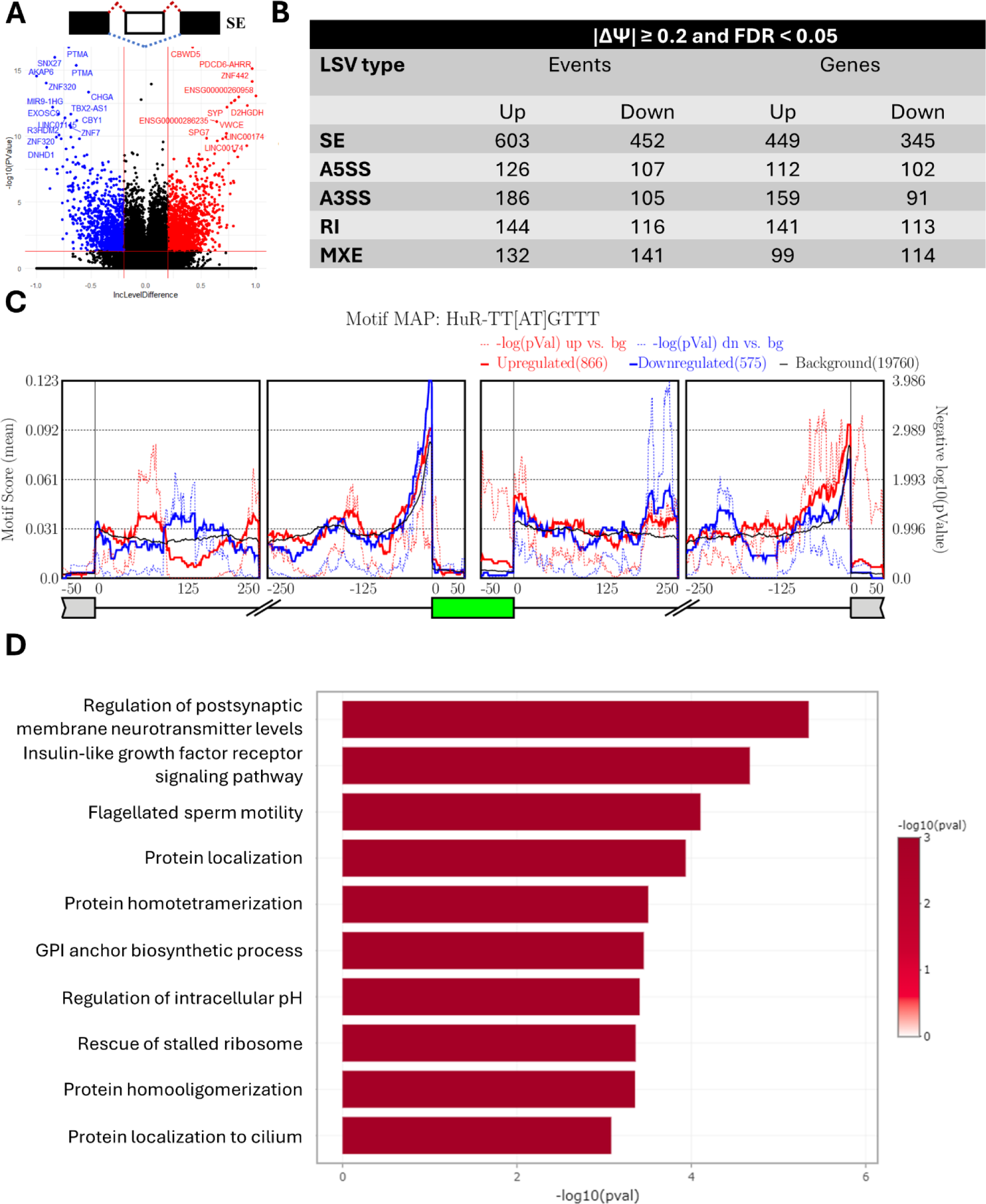
HuR KD induces transcriptome-wide alternative splicing changes. **A:** Volcano plot of skipped exon events after HuR KD. **B:** Table showing the number of significant events and differentially spliced genes in each LSV program after HuR KD. **C:** Motif-map showing the enrichment of HuR motif in the downregulated SE events after HuR KD. **D:** ORA analysis showing the enrichment of GOBP terms in the downregulated SE genes after HuR KD.

In summary, HuR is downregulated during neuronal differentiation to promote exon skipping of multiple genes involved in the differentiation process and mature neuronal function.

## Discussion

Alternative splicing has long been recognized as an important mechanism involved in neural development and plasticity (30,31). Nonetheless, most of the attempts to understand AS regulation during neural differentiation and development focused on a single interactor or RBP, with few studies attempting transcriptome-wide analysis of AS in neural models (32,33). Here, we used SH-SY5Y neural differentiation model to study AS. SH-SY5Y was particularly intriguing given its heavy use is studying neurodegenerative diseases such as Parkinson’s disease (PD) and its acceptability as a model for neural functional analysis in the neuroscience community (2,13,15,34). While several studies have attempted transcriptomics analysis of this model of neural differentiation, the epi and post-transcriptional landscape of differentiated mature neurons from SH-SY5Y cells remain unknown. Previously, we characterized mRNA stability changes, and identified how such changes in mRNA stability drive neural differentiation of SH-SY5Y cells and impact their response to oxidative stress (6). Here, we expanded our work to analyze AS transcriptome-wide, as well as identifying its regulators. Our analysis revealed transcriptome-wide changes in AS, dominated by alterations in exon skipping, during neural differentiation. These changes were driven by various RBPs, of which, we selected PTB and HuR, as the top RBP motifs enriched in SE events, for downstream validation. PTB and HuR impacted neural differentiation and phenotypes by driving AS of many mRNAs that are related to neuronal function and homeostasis.

Approximately 95% of human multi-exon genes undergo alternative splicing, which can lead to the generation of various proteoforms that have different functions or subcellular localizations (31). Such plasticity of the transcriptome is essential for cellular specialization, functioning, and development (31,32,35). The regulation of mRNA alternative splicing is a complex process that entails dynamic interplay between multiple factors such as mRNA modifications, most famously N6-methyladenosine or m^6^A (36), and RBPs acting as splice activators or repressors via interacting with spliceosome components (30,35). While alternative splicing has been long recognized for its roles in modulating neuronal maturation, migration, synapse formation, and various functions (31–33,35), most early transcriptomic studies lacked enough depth to allow for proper characterization of global AS events as well as their regulatory frameworks (32). Despite these limitations, many important temporal events were recognized to play an important role in brain development. For example, dynamic changes in RBPs drive neuronal maturation and differentiation during development, and alteration of such RBPs dysregulates neuronal differentiation (reviewed in (31,35,37)). Recent advances and improvements in the sequencing technologies as well as bioinformatics tools allowed unprecedented insights into the temporal AS events occurring during brain development, down to the single cell level (30,33). With these developments in mind, we were motivated to bring such insights into one of the most important and widely used cellular models used to study neurodegenerative diseases and neural function, the SH-SY5Y neural differentiation model. Despite its ubiquity, only transcriptomic profiling of this model was previously focused on (6). While we previously reported the changes in mRNA stability and translation during SH-SY5Y differentiation, as well as their regulation via RBPs such as SAMD4A (6), there remains a large gap in our knowledge of this model. Considering that a simple search on PubMed using the keywords “shsy5y” and “Parkinson” yielded >2,400 results, it becomes clear why we need to understand this cellular model in detail. While direct parallels between neurons developed from SH-SY5Y neuroblastoma cells and cortical neurons should not be easily made, we observed many similarities between both that indicate the robustness of the SH-SY5Y model in studying neural AS. For example, a dominance of exon skipping events on the AS landscape (30,38) and the regulation of AS by RBPs known for their role in neural differentiation and function such as PTB/PTBP1 (39), HuR/ELAVL1 (27), and SRP20 (40).

The PTB family consists of 3 members; PTB/PTBP1, which is expressed in most cells, PTBP2 (nPTB) which is expressed in neurons, and PTBP3/ROD1, which is expressed in immune cells (39). PTB/PTBP1 is expressed only during the embryonic stage in neurons (39). PTB/PTBP1 suppresses the inclusion of exon 10 of PTBP2, leading to its degradation via non-sense mediated decay (31,41) During development PTB/PTBP1 is suppressed by the neural specific miR-124 (41). This suppression of PTB/PTBP1 leads to the inclusion of exon 10 of PTBP2 and the activation of its protein expression, driving the neurons towards differentiation (31). Once neurons developed synapses, PTBP2 is in turn downregulated (31). This orchestrated and complex sequence of events drive the neuronal development via regulating various temporal changes at the level of AS (30). In addition, the links between PTB downregulation and neuronal differentiation were explored as a potential mechanism to induce direct neuronal reprogramming of astrocytes, that could be used for neurological diseases therapies (23). However, such claims did not hold to scrutiny (42). In our analysis, we observed downregulation of PTB/PTBP1 during neuronal differentiation at the protein level. This downregulation was associated with global activation of exon skipping programs in multiple mRNAs, a feature that was replicated in PTB KD cells. In addition, PTB was found to drive the AS of genes involved in various neuronal maturation and differentiation related pathways, in agreement with previous literature.

ELAVL1/HuR is known to play roles in neural stem cells differentiation and neural crest development (27,43). It is also known to play important roles in neurodegenerative disorders (44). Nonetheless, its role in neuronal differentiation is less defined than that of PTB. Here, we show that during neuronal differentiation, HuR levels are downregulated, in tandem with reduced exon skipping of HuR bound mRNAs. Pathways related to HuR were also shown to be related to neuronal functioning and development as we show herein.

An important reason for the need to develop our understanding of the posttranscriptional changes during neuronal development that dictate neuronal phenotype and function, is the close ties between such phenomena and neurodegenerative diseases (44,45). AS is recognized as one of the core phenomena associated with various neurodegenerative diseases such as ALS, Alzheimer’s and more (45). Given that the SH-SY5Y model is used to study neurodegenerative disease, particularly Parkinson’s disease (PD) via respiratory complex I inhibition, we added another layer to our analysis by interrogating the impact of the studied RBPs on neuronal sensitivity to stresses. We have shown previously that mature neurons are more sensitive to various specific stressors (6). This is in line with the known sensitivity of neurons to oxidative stress, that is a core pathophysiological mechanism in neurodegenerative and neurological diseases (46,47). Here we show that downregulation of either PTB or HuR sensitizes SH-SY5Y cells to mitochondrial stress and ferroptosis. Given that we also observed changes in the programmable cell death activated during stress, akin to our previous observations in mature neurons (6), we postulate that such sensitivity is secondary to pushing the cells towards a more differentiated phenotype.

In this study, we present a wealth of data on an important model for studying neural differentiation and neurodegeneration. However, the study has certain limitations that should be considered. While we opted for a more computational and bioinformatics driven approach, downstream effects of PTB and HuR still need to be further characterized. In addition, how much of the findings from this model would correlate with actual changes in brain neurons should be also considered. While our results agree with previous work done using brains or primary neurons, we do not believe it is a 100% match.

In conclusion, we present a comprehensive transcriptome-wide analysis of alternative mRNA splicing events in various LSV programs during neuronal differentiation of SH-SY5Y neuroblastoma cells. We show that RBPs regulate various LSV programs, and we further validate two of these RBPs; HuR and PTB. While the AS events were mostly of exon skipping types, we believe further work should aim to characterize the other reported LSV subtypes such as intron retention or alternative exon splicing. In addition, proteogenomic approaches should elucidate the true impact of AS on the proteome. Understanding these programs, and their implications in the context of neural development and function, is critical for our understanding of brain function, development, and diseases.

## Methods

### Cell culture

Human SH-SY5Y neuroblastoma cells obtained from ATCC (Cat# CRL-2266) were cultured in Eagle’s Minimum Essential Medium (EMEM; ATCC, Cat# 30-2003) and Ham’s F-12 Nutrient Mixture (F12; Gibco, Cat# 11765054) 1:1 containing 10% heat-inactivated Fetal Bovine Serum (FBS; Corning, Cat# 27419002), at 37℃ and 5% CO2. No antibiotics were added to the growth media.

### Differentiation protocol

The neuronal differentiation protocol for SH-SY5Y cells was conducted as previously reported (6). The differentiation protocol consisted of two stages: a 4-day pre-differentiation step in EMEM/F-12 supplemented with 10% FBS and 10uM RA (Sigma-Aldrich, Cat# R2625), and a subsequent 6-day differentiation step in serum-free medium containing 50 ng/ml human brain derived neurotrophic factor (BDNF) (Sigma-Aldrich, Cat# B3795). Cells were seeded at an initial density of 2 × 10^4^ cells/cm^2^ in 24-well plate coated with Type I collagen (Corning, Cat# 3524). Media were routinely changed every 2-3 days.

### Western blot

Cells seeded in 6 well plates were homogenized in T-PER reagent (ThermoFisher, Cat# 78510), 1% Triton X-100, 1x complete protease inhibitor (Roche, Cat# 4693116001) and 1x phosphatase inhibitor (Roche, Cat# 4906845001) on ice for 30 min. The cell lysate was then sonicated and centrifuged at 16,000g for 15min and the supernatants were extracted for protein concentration measurement by Pierce BCA protein assay (ThermoFisher, Cat# 23227). 20-30μg protein samples were loaded into 4-20% Mini-PROTEIN TGX Precast Protein Gels (Bio-Rad, Cat# 4561096), and then were transferred to 0.2µm nitrocellulose membranes (Bio-Rad, Cat# 1704158). The membranes were blocked by 5% skim milk power (Nacalai Tesque, Cat# 31149-75) in 1x Phosphate buffered saline with Tween (PBS-T, Sigma-Aldrich, Cat# 524653), followed by overnight incubation with primary antibodies at 4 ℃. After incubation, the membranes were washed three times with PBS-T and then incubated with secondary antibody at room temperature for 1 hour. The membranes were washed three times with PBS-T again and then incubated with Pierce™ ECL Western Blotting Substrate (ThermoFisher, Cat# 32106). The protein bands were visualized by ChemiDoc MP (BioRad). Digital images were processed and analyzed using Image J.

### Primary antibodies used

Nestin (Invitrogen, Cat#MA1-110)

MAP2 (Sigma, Cat#M1406)

TUJ (Cell Signaling, Cat#2128)

SYN (Invitrogen, Cat#MA5-14532)

PTB/PTBP1 (Invitrogen, Cat#32-5000)

HuR (Invitrogen, Cat#MA1-167)

beta-actin (Cell Signaling, Cat#4970)

### Secondary antibodies used

Rabbit IgG HRP (Cell Signaling, Cat#7074)

Mouse IgG HRP (Cell Signaling, Cat#7076)

### Stress response assay

Five different stressors from three categories of stress were selected: sodium arsenite (AS) to induce nonspecific oxidative stress, Antimycin A (AntiA; respiratory complex III inhibitor) and Rotenone (Rot; respiratory complex I inhibitor) as mitochondria stressors, and Erastin (Er) and 1S, 3R-RSL3 (RSL) to induce ferroptosis stress. Each of these stressors has been reported to play a significant role in neuronal damage and the pathophysiology of neurodegenerative disorders(Bhat et al., 2015). By combining the treatment of different stress inhibitors: Ferrostatin-1 (Fer-1, ferroptosis inhibitor), Necrostatin-1 (Nec-1, necroptosis inhibitor), Ac-YVAD-cmk (YVAD, Pyroptosis inhibitor) and Ac-DVED-CHO (DEVD, apoptosis inhibitor), the main program of cell death induced by stressors could be identified.

After gene KD, cells were seeded on flat bottom 96-well plates with collagen coated at 10,000 cells/well. Cells were pretreated with four stress inhibitors: 20μM Fer-1 (Sigma-Aldrich, Cat# SML0583), 40μM Nec-1 (Sigma-Aldrich, Cat# N9037), 10μM YVAD (Sigma-Aldrich, Cat# SML0429) and 10μM DEVD (Sigma-Aldrich, Cat# A0835) for 1 hour. Later, five different stressors: 4μM AntiA (Sigma-Aldrich, Cat# N8674), 0.2μM Rot (Sigma-Aldrich, Cat# 557368), 12μM AS (Sigma-Aldrich, Cat# S7400), 20μM Er (Sigma-Aldrich, Cat# E7781) and 2μM RSL (Sigma-Aldrich, Cat# SML2234) were added to inhibitor-pretreated-groups and stressor-only-groups for 24 hours. The control groups were treated with the same amount of growth media.

Cells were then incubated with WST-8 (Nacalai tesque, Cat# 07553-44) for 4 hours at 37℃. The absorbance of formazan was measured at 450nm by Spectra Max microplate reader (Molecular Devices Spectramax 190). Each treatment had 6 replicate wells and was repeated twice (a total of 12 biological replicates). Cell viability was expressed as a percentage of the control.

### RNA isolation and quality control

Cells were lysed in QIAzol Lysis Reagent (QIAGEN, Cat# 79306) for RNA extraction. RNA was extracted using the miRNeasy Kit (QIAGEN Cat# 217004) with a DNase digestion step following the manufacturer’s instructions. RNA purity and concentration were examined by Nanodrop (Thermo Fisher Scientific; Catalog# ND-ONE-W), and RNA integrity number (RIN) was assessed using RNA 6000 Nano Kit (Agilent, Cat# 5067-1511) on Agilent Bioanalyzer 2100. Samples with RNA integrity number ≥ 9 were used.

### RNA sequencing (RNA-seq)

RNA-seq libraries were prepared from three biological replicates of Mock or KD cells using NEBNext Poly(A) mRNA magnetic isolation module (NEB, Cat# E7490) for mRNA enrichment and Ultra II directional RNA Library Prep Kit (NEB, Cat# E7760) following the manufacturer’s instruction. The quality of libraires was assessed by Agilent DNA 1000 kit (Agilent, Cat# 5067-1504)) on the Agilent Bioanalyzer 2100. The concentration of libraries was determined using NEBNext library Quant kit for Illumina (NEB, Cat# E7630). Libraries were then pooled and sequenced by Macrogen on Ilumina Hiseq X-ten platform. The sequencing was performed with 150 base-pair pair-end reads (150bp × 2) and the target depth was 50 million per sample.

### Processing of the sequencing data

Quality control for Raw Fastq was performed using FastQC. Raw reads were then trimmed with Trimmomatic (48) and adaptor sequences and low-quality read removed. Reads were aligned to the human reference genome hg38 (GRCh38.p13) using the splice aware aligner HISAT2 (49). Around 84.5%-87.5% of read pairs were uniquely mapped to the hg38 genome. Mapped reads BAM file were then counted to gene features by FeatureCounts (50) with standard settings. Cutoff for significant gene expression was FDR < 0.05 and |Log_2_ FC| ≥ 1. Local splice variants (LSV) analysis was conducted using rMATs (16) and motif analysis of RBPs was conducted using rMAPs2 (22). Cutoff value for statistically significant splice event was FDR < 0.05 and |ΔΨ| ≥ 0.02. Pre ranked gene set enrichment analysis was conducted using eVITTA toolbox (51). Overrepresentation analysis (ORA) was conducted using eVITTA or Metascape (52).

## Supporting information

Suppmenetary tables

## Data visualization

Volcano plots were created using R language programming. Heatmaps were created using Morpheus software (https://software.broadinstitute.org/morpheus/)

## Short Hairpin RNA design (shRNA) and knockdown experiment

Short hairpin RNA (shRNA)-encoding pairs of oligonucleotides with targeting sequences to PTB and HuR mRNA was designed as follows:

PTB:

5’TGCTTCTGCAGCAAACGGAAATTTCAAGAGAATTTCCGTTTGCTGCAGAAGCTTTT TTC

HuR:

5’TCGAGCTCAGAGGTGATCAAAGTTCAAGAGACTTTGATCACCTCTGAGCTCGTTTT TTC

Mock:

5’TGAAATACTCAGCAGATCATTATTCAAGAGATAATGATCTGCTGAGTATTTCTTTT TTC-3’

Each sequence was cloned into the corresponding sites of pLB vector (Addgene, Cat#11619). Lentiviruses were generated by co-transfection of Lenti-X 293T (Takara, Cat# 632180) cells with three plasmids: a lentiviral vector plasmid, pMD2.G (expressing envelop protein, FASMAC) and psPAX2 (expressing packaging proteins, FASMAC). Media were changed 16 h after transfection, and the supernatants were harvested 48 h after transfection. Cell debris in the media was removed by 0.45 µm filtration following centrifugation at 1500 g for 10 min. Viral particles were collected twice 48 h and 72 h post-transfection. For infection, lentivirus particles were added to each well of a six-well plate containing 7.5 × 105 cells. Cells were incubated with lentivirus and 4 µg/ml polybrene (Sigma, Cat#TR-1003) for 12 h. The expression of Green Fluorescent Protein (GFP) was checked under the immunofluorescence microscopy. The transfection efficiency of cells was set to 100% based on fluorescence distribution and western blot was used to evaluate the knockdown efficiency.

### Neurite Growth Rate Analysis

The analysis of neurite growth rate was conducted using the images obtained from the microscope (Leica DMi1). Image J was utilized to track the length of neurites (Ridge Detection) and the surface area of cells, including both neurites and cell bodies. The ratio of neurite length to cell surface area was calculated as the percentage of neurite outgrowth activity.

### Statistical Analysis

Statistical tests were performed with GraphPad Prism 7 software. The values were presented as mean ± standard deviation (SD) of at least 3 biological replicates or as indicated. For each dataset, the Shapiro-Wilk normality test was applied to determine if the data had a normal distribution. If the data passed the normality test, the parametric test was used, otherwise the non-parametric test was used. Analysis of western blot data was performed with unpaired Student t-test (two-tailed). For the analysis of stress response assay and neurite growth rate, two-way analysis of variance (ANOVA) with Bonferroni post-hoc test was performed. The statistical significance. The statistical significance was set at P < 0.05 (*P < 0.05, **P < 0.01, ***P < 0.001 and ****P < 0.0001).

## Data availability

RNA sequencing data of PTB and HuR KD was deposited in the sequence read archive (PRJNA1001516). The RNA-seq and Ribo-seq data of differentiated neurons was retrieved from previous publication (6) and is available through the sequence read archive (PRJNA1001994 and PRJNA779467).

## Author contributions

**YZ:** Experimentation. Protocol design. Data analysis. Manuscript revision. Funding. **SR:** Conception and study design. Data analysis. Writing. Supervision. Funding **KN:** Critically revised the manuscript. Supervision.

## Acknowledgements

The authors report no conflict of interest nor there are any ethical adherences regarding this work.

