## Supplementary figures for "Transcriptome-wide alternative mRNA splicing analysis reveals post-transcriptional regulation of neuronal differentiation"

**Supplementary figure 1: Analysis of the splicing machinery components after neuronal differentiation. A:** Enrichment of Spliceosome pathway at the level of gene expression after neuronal differentiation. **B:** Kegg native view showing the expression of various spliceosome components in the RNA-seq dataset. **C:** Enrichment of Spliceosome pathway at the level of mRNA translation after neuronal differentiation. **D:** Kegg native view showing the expression of various spliceosome components in the Ribo-seq dataset. Volcano plot showing the differential expression of various RBPs involved in alternative splicing after neuronal differentiation from the RNA-seq **(E)** and Ribo-seq **(F)** datasets.

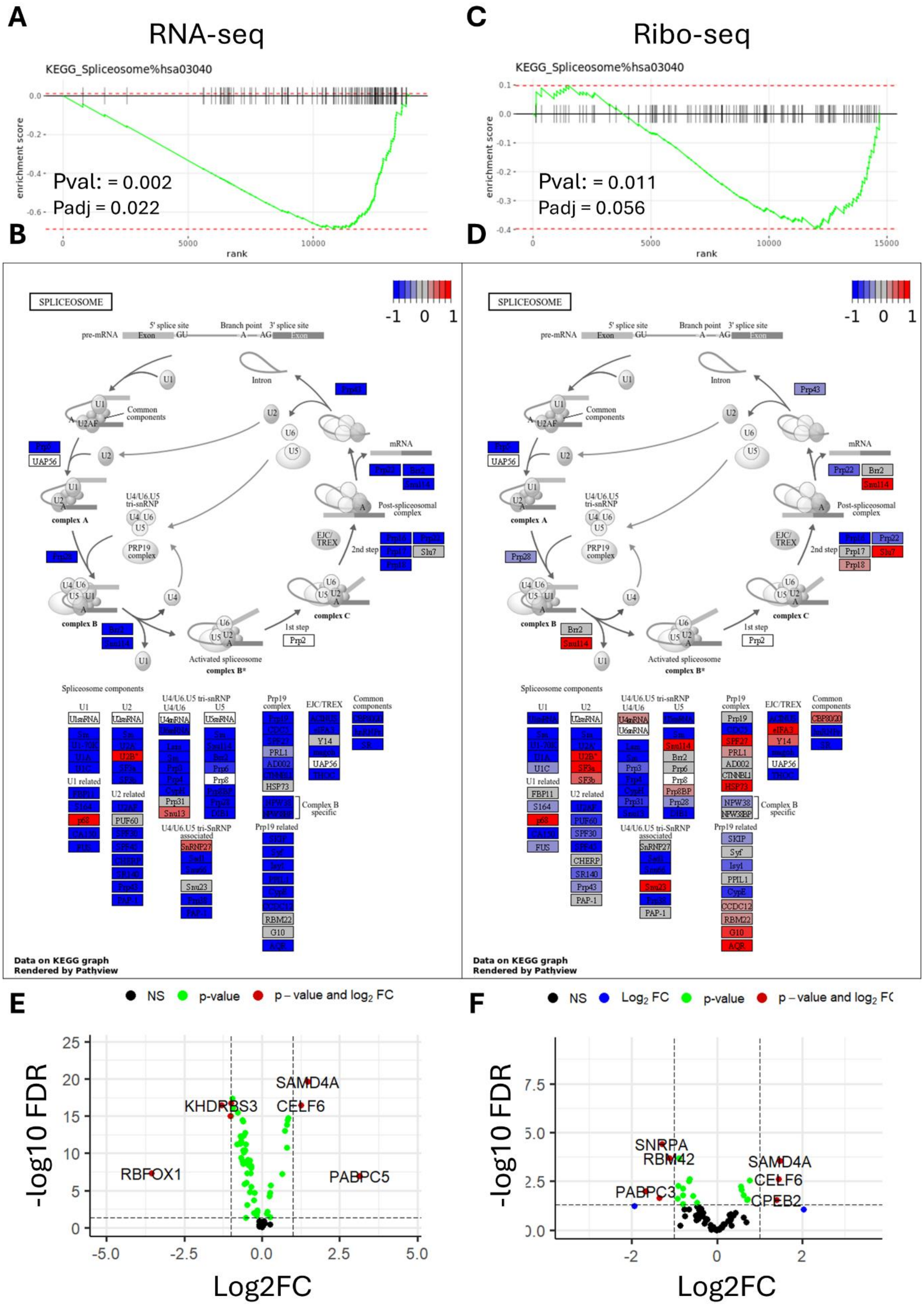
A
RNA-seq
KEGG_Spliceosome%hsa03040
enrichment score
Pval: = 0.002
Padj = 0.022
rank
C
Ribo-seq
KEGG_Spliceosome%hsa03040
enrichment score
Pval: = 0.011
Padj = 0.056
rank
B
D
SPLICEOSOME
-1 0 1
pre-mRNA
Exon
5' splice site
Branch point
3' splice site
Intron
Common components
U2AF
UAP56
complex A
U4/U6.U5 tri-snRNP
PRP19 complex
complex B
Activated spliceosome complex B*
1st step
Prp2
complex C
2nd step
Slu7
EJC/ TREX
Post-spliceosomal complex
mRNA
Spliceosome components
U1
U2
U4/U6
U5
Prp19 complex
EJC/TREX
Common components
U1 related
U2 related
U4/U6.U5 tri-SnRNP associated
Prp19 related
Complex B specific
Data on KEGG graph
Rendered by Pathview
E
NS
p-value
p – value and log2 FC
-log10 FDR
Log2FC
SAMD4A
KHDRBS3
CELF6
RBFOX1
PABPC5
F
NS
Log2 FC
p-value
-log10 FDR
Log2FC
SNRPA
RBM42
SAMD4A
PABPC3
CELF6
CPEB2

**Supplementary figure 2:** Motif-maps of HuR motifs enrichment (continuation of Figure 2D)

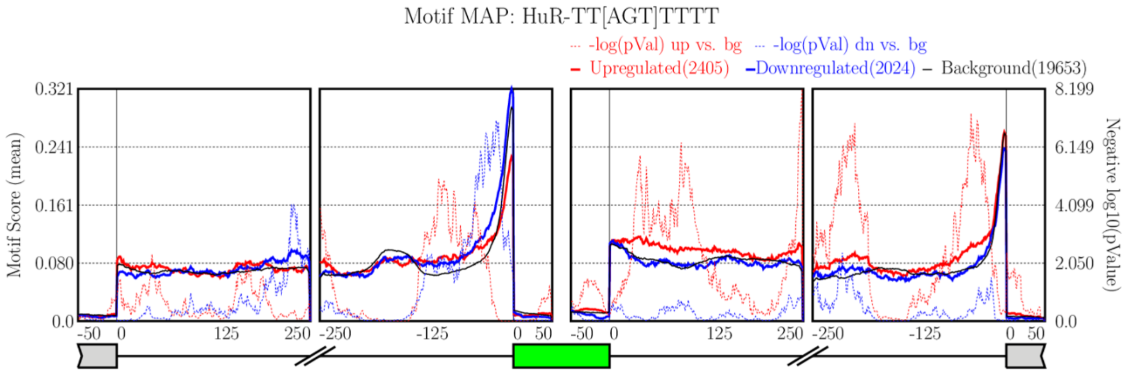
Motif MAP: HuR-TT[AGT]TTTT
-log(pVal) up vs. bg
-log(pVal) dn vs. bg
Upregulated(2405)
Downregulated(2024)
Background(19653)
Motif Score (mean)
Negative log10(pValue)
0.321
0.241
0.161
0.080
0.0
8.199
6.149
4.099
2.050
0.0
-50 0 125 250 -250 -125 0 50 -50 0 125 250 -250 -125 0 50

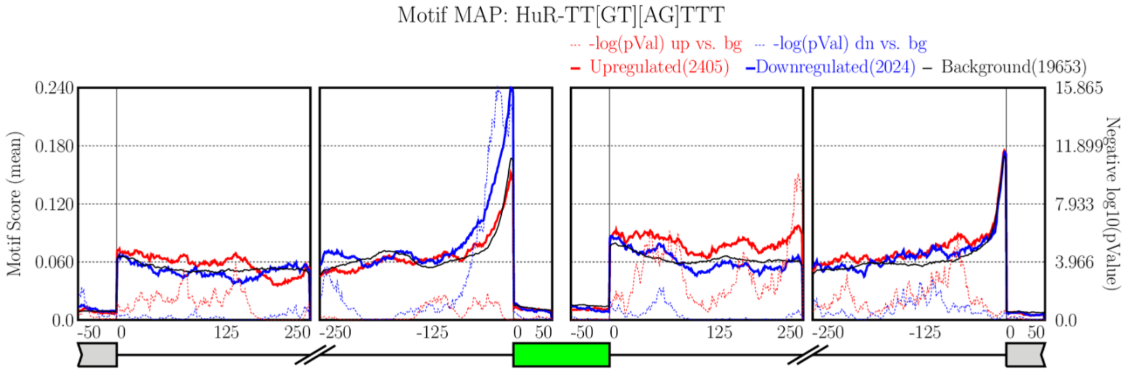
Motif MAP: HuR-TT[GT][AG]TTT
-log(pVal) up vs. bg
-log(pVal) dn vs. bg
Upregulated(2405)
Downregulated(2024)
Background(19653)
Motif Score (mean)
Negative log10(pValue)
0.240
0.180
0.120
0.060
0.0
15.865
11.899
7.933
3.966
0.0
-50 0 125 250 -250 -125 0 50 -50 0 125 250 -250 -125 0 50

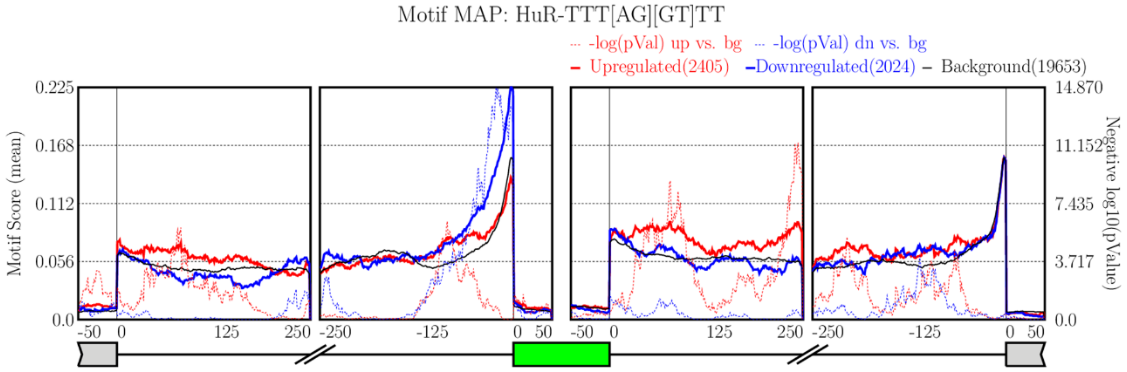
Motif MAP: HuR-TTT[AG][GT]TT
-log(pVal) up vs. bg
-log(pVal) dn vs. bg
Upregulated(2405)
Downregulated(2024)
Background(19653)
Motif Score (mean)
Negative log10(pValue)
0.225
0.168
0.112
0.056
0.0
14.870
11.152
7.435
3.717
0.0
-50 0 125 250 -250 -125 0 50 -50 0 125 250 -250 -125 0 50

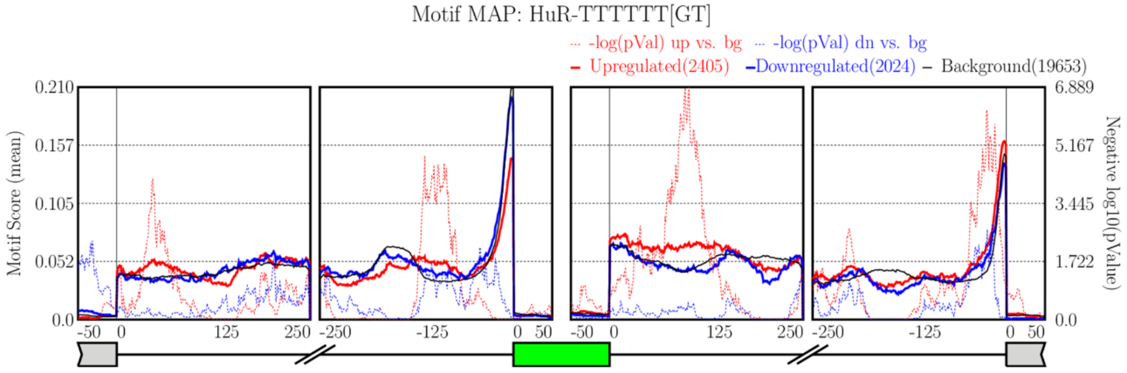
Motif MAP: HuR-TTTTTT[GT]
-log(pVal) up vs. bg
-log(pVal) dn vs. bg
Upregulated(2405)
Downregulated(2024)
Background(19653)
Motif Score (mean)
Negative log10(pValue)
0.210
0.157
0.105
0.052
0.0
6.889
5.167
3.445
1.722
0.0
-50 0 125 250 -250 -125 0 50 -50 0 125 250 -250 -125 0 50

**Supplementary figure 3: Analysis of RBP motifs involved in intron retention events. A:** Heatmaps of motifs enriched in the upregulated RI events. **B:** Heatmaps of motifs enriched in the downregulated RI events. **C-E:** Motif-map of the enrichment of the top enriched RBPs motifs.

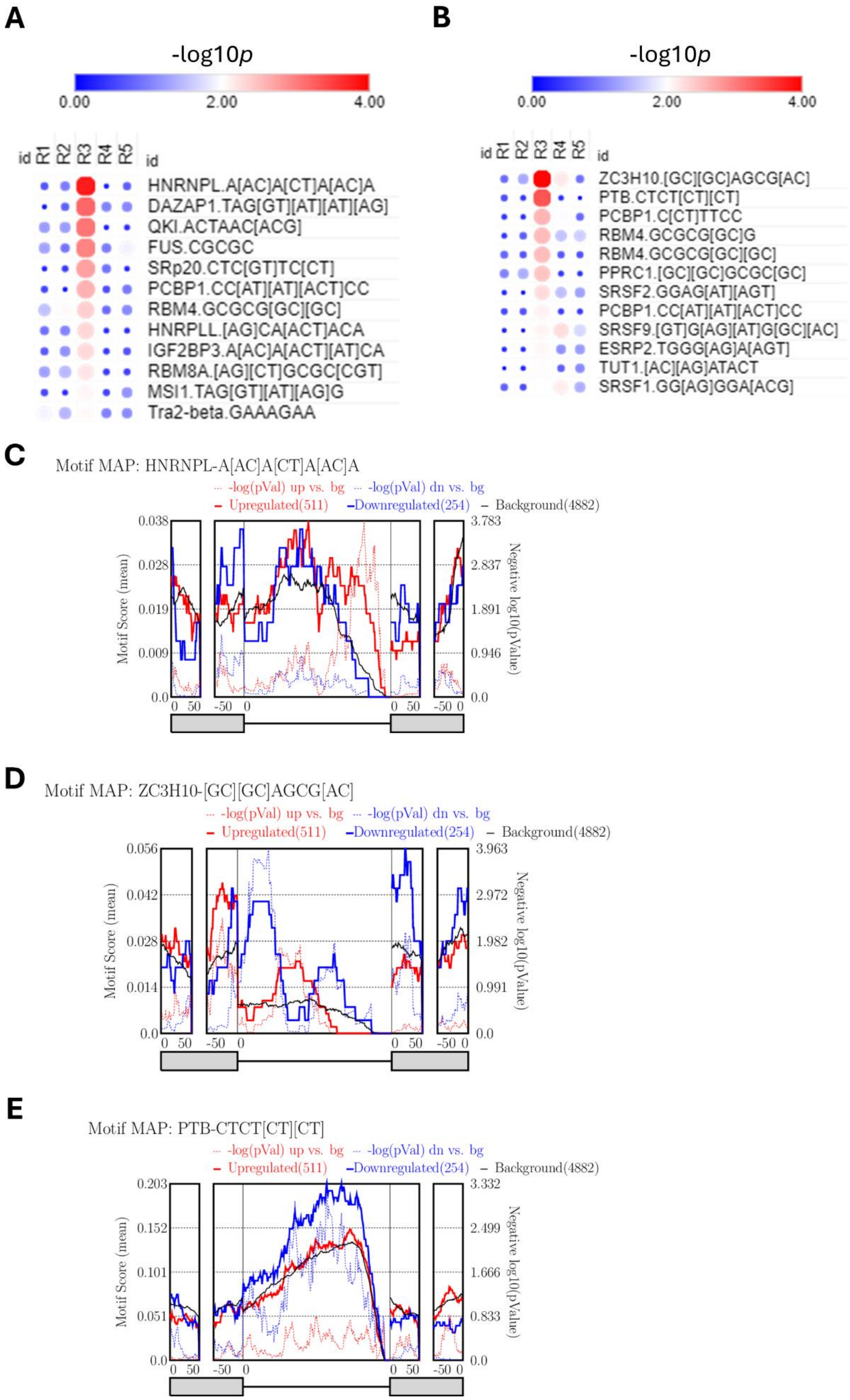

A
-log10p
0.00
2.00
4.00
id R1 R2 R3 R4 R5 id
HNRNPL.A[AC]A[CT]A[AC]A
DAZAP1.TAG[GT][AT][AT][AG]
QKI.ACTAAC[ACG]
FUS.CGCGC
SRp20.CTC[GT]TC[CT]
PCBP1.CC[AT][AT][ACT]CC
RBM4.GCGCG[GC][GC]
HNRPLL.[AG]CA[ACT]ACA
IGF2BP3.A[AC]A[ACT][AT]CA
RBM8A.[AG][CT]GCGC[CGT]
MSI1.TAG[GT][AT][AG]G
Tra2-beta.GAAAGAA
B
-log10p
0.00
2.00
4.00
id R1 R2 R3 R4 R5 id
ZC3H10.[GC][GC]AGCG[AC]
PTB.CTCT[CT][CT]
PCBP1.C[CT]TTCC
RBM4.GCGCG[GC]G
RBM4.GCGCG[GC][GC]
PPRC1.[GC][GC]GCGC[GC]
SRSF2.GGAG[AT][AGT]
PCBP1.CC[AT][AT][ACT]CC
SRSF9.[GT]G[AG][AT]G[GC][AC]
ESRP2.TGGG[AG]A[AGT]
TUT1.[AC][AG]ATACT
SRSF1.GG[AG]GGA[ACG]
C
Motif MAP: HNRNPL-A[AC]A[CT]A[AC]A
-log(pVal) up vs. bg
-log(pVal) dn vs. bg
Upregulated(511)
Downregulated(254)
Background(4882)
Motif Score (mean)
Negative log10(pValue)
0.038
0.028
0.019
0.009
0.0
3.783
2.837
1.891
0.946
0.0
0 50
-50 0
0 50
-50 0
D
Motif MAP: ZC3H10-[GC][GC]AGCG[AC]
-log(pVal) up vs. bg
-log(pVal) dn vs. bg
Upregulated(511)
Downregulated(254)
Background(4882)
Motif Score (mean)
Negative log10(pValue)
0.056
0.042
0.028
0.014
0.0
3.963
2.972
1.982
0.991
0.0
0 50
-50 0
0 50
-50 0
E
Motif MAP: PTB-CTCT[CT][CT]
-log(pVal) up vs. bg
-log(pVal) dn vs. bg
Upregulated(511)
Downregulated(254)
Background(4882)
Motif Score (mean)
Negative log10(pValue)
0.203
0.152
0.101
0.051
0.0
3.332
2.499
1.666
0.833
0.0
0 50
-50 0
0 50
-50 0

**Supplementary figure 4: Analysis of RBP motifs involved in 5' alternative exon splice site (A5SS) events. A:** Heatmaps of motifs enriched in the upregulated A5SS events. **B:** Heatmaps of motifs enriched in the downregulated A5SS events. **C-D:** Motif-map of the enrichment of the top enriched RBPs motifs.

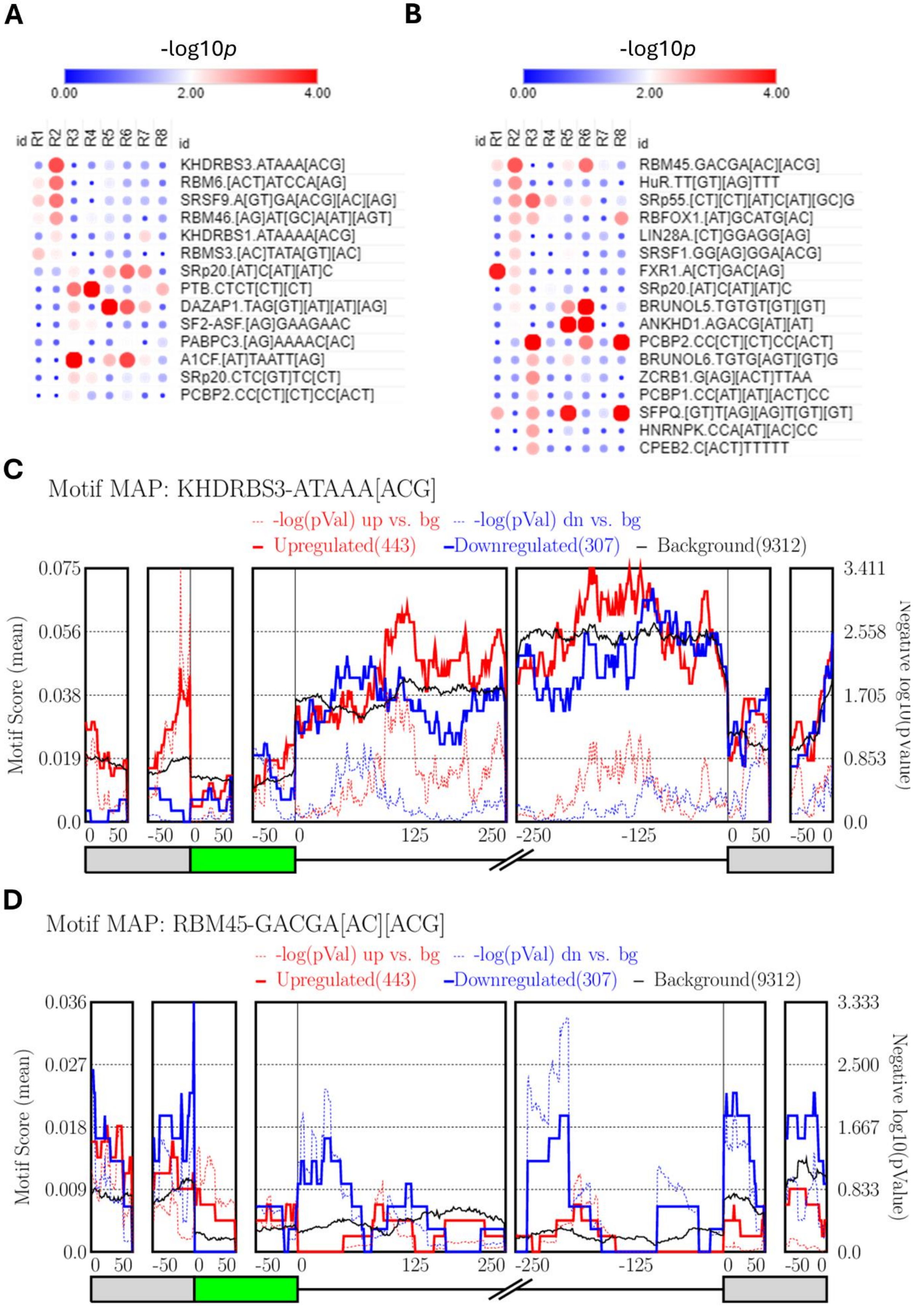

A
B
-log10p
0.00
2.00
4.00
id R1 R2 R3 R4 R5 R6 R7 R8 id
KHDRBS3.ATAAA[ACG]
RBM6.[ACT]ATCCA[AG]
SRSF9.A[GT]GA[ACG][AC][AG]
RBM46.[AG]AT[GC]A[AT][AGT]
KHDRBS1.ATAAAA[ACG]
RBMS3.[AC]TATA[GT][AC]
SRp20.[AT]C[AT][AT]C
PTB.CTCT[CT][CT]
DAZAP1.TAG[GT][AT][AT][AG]
SF2-ASF.[AG]GAAGAAC
PABPC3.[AG]AAAAC[AC]
A1CF.[AT]TAATT[AG]
SRp20.CTC[GT]TC[CT]
PCBP2.CC[CT][CT]CC[ACT]
RBM45.GACGA[AC][ACG]
HuR.TT[GT][AG]TTT
SRp55.[CT][CT][AT]C[AT][GC]G
RBFOX1.[AT]GCATG[AC]
LIN28A.[CT]GGAGG[AG]
SRSF1.GG[AG]GGA[ACG]
FXR1.A[CT]GAC[AG]
SRp20.[AT]C[AT][AT]C
BRUNOL5.TGTGT[GT][GT]
ANKHD1.AGACG[AT][AT]
PCBP2.CC[CT][CT]CC[ACT]
BRUNOL6.TGTG[AGT][GT]G
ZCRB1.G[AG][ACT]TTAA
PCBP1.CC[AT][AT][ACT]CC
SFPQ.[GT]T[AG][AG]T[GT][GT]
HNRNPK.CCA[AT][AC]CC
CPEB2.C[ACT]TTTTT
C
Motif MAP: KHDRBS3-ATAAA[ACG]
-log(pVal) up vs. bg
-log(pVal) dn vs. bg
Upregulated(443)
Downregulated(307)
Background(9312)
Motif Score (mean)
Negative log10(pValue)
0.075
0.056
0.038
0.019
0.0
3.411
2.558
1.705
0.853
0.0
0 50
-50 0 50
-50 0 125 250
-250 -125 0 50
-50 0
D
Motif MAP: RBM45-GACGA[AC][ACG]
-log(pVal) up vs. bg
-log(pVal) dn vs. bg
Upregulated(443)
Downregulated(307)
Background(9312)
Motif Score (mean)
Negative log10(pValue)
0.036
0.027
0.018
0.009
0.0
3.333
2.500
1.667
0.833
0.0
0 50
-50 0 50
-50 0 125 250
-250 -125 0 50
-50 0

**Supplementary figure 5: Analysis of RBP motifs involved in 3' alternative exon splice site (A3SS) events. A:** Heatmaps of motifs enriched in the upregulated A3SS events. **B:** Heatmaps of motifs enriched in the downregulated A3SS events. **C-D:** Motif-map of the enrichment of the top enriched RBPs motifs.

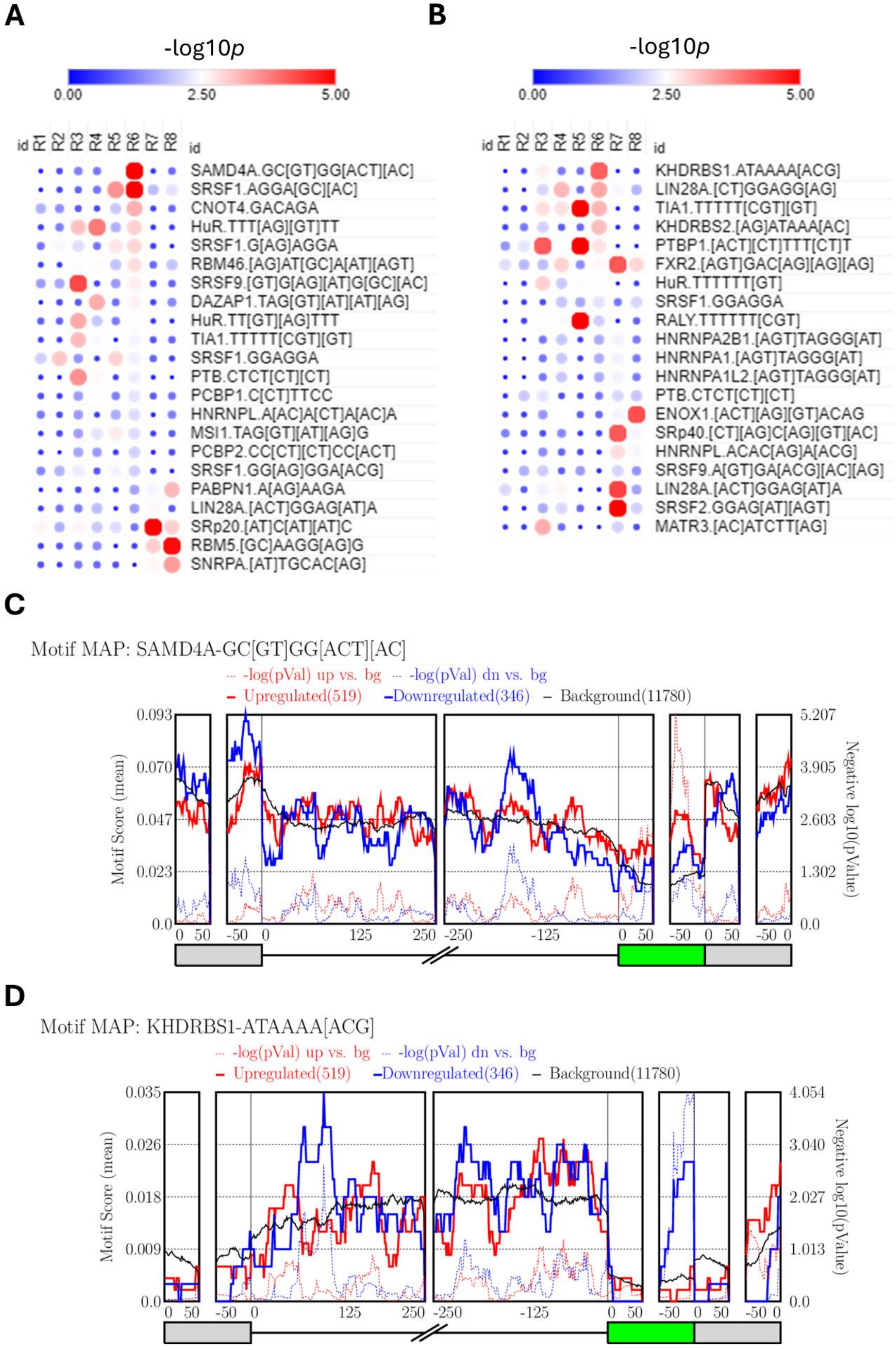
A
-log10p
0.00
2.50
5.00
id R1 R2 R3 R4 R5 R6 R7 R8 id
SAMD4A.GC[GT]GG[ACT][AC]
SRSF1.AGGA[GC][AC]
CNOT4.GACAGA
HuR.TTT[AG][GT]TT
SRSF1.G[AG]AGGA
RBM46.[AG]AT[GC]A[AT][AGT]
SRSF9.[GT]G[AG][AT]G[GC][AC]
DAZAP1.TAG[GT][AT][AT][AG]
HuR.TT[GT][AG]TTT
TIA1.TTTTT[CGT][GT]
SRSF1.GGAGGA
PTB.CTCT[CT][CT]
PCBP1.C[CT]TTCC
HNRNPL.A[AC]A[CT]A[AC]A
MSI1.TAG[GT][AT][AG]G
PCBP2.CC[CT][CT]CC[ACT]
SRSF1.GG[AG]GGA[ACG]
PABPN1.A[AG]AAGA
LIN28A.[ACT]GGAG[AT]A
SRp20.[AT]C[AT][AT]C
RBM5.[GC]AAGG[AG]G
SNRPA.[AT]TGCAC[AG]
B
-log10p
0.00
2.50
5.00
id R1 R2 R3 R4 R5 R6 R7 R8 id
KHDRBS1.ATAAAA[ACG]
LIN28A.[CT]GGAGG[AG]
TIA1.TTTTT[CGT][GT]
KHDRBS2.[AG]ATAAA[AC]
PTBP1.[ACT][CT]TTT[CT]T
FXR2.[AGT]GAC[AG][AG][AG]
HuR.TTTTTT[GT]
SRSF1.GGAGGA
RALY.TTTTTT[CGT]
HNRNPA2B1.[AGT]TAGGG[AT]
HNRNPA1.[AGT]TAGGG[AT]
HNRNPA1L2.[AGT]TAGGG[AT]
PTB.CTCT[CT][CT]
ENOX1.[ACT][AG][GT]ACAG
SRp40.[CT][AG]C[AG][GT][AC]
HNRNPL.ACAC[AG]A[ACG]
SRSF9.A[GT]GA[ACG][AC][AG]
LIN28A.[ACT]GGAG[AT]A
SRSF2.GGAG[AT][AGT]
MATR3.[AC]ATCTT[AG]
C
Motif MAP: SAMD4A-GC[GT]GG[ACT][AC]
-log(pVal) up vs. bg
-log(pVal) dn vs. bg
Upregulated(519)
Downregulated(346)
Background(11780)
Motif Score (mean)
Negative log10(pValue)
0.093
0.070
0.047
0.023
0.0
5.207
3.905
2.603
1.302
0.0
0 50
-50 0 125 250
-250 -125 0 50
-50 0 50
-50 0
D
Motif MAP: KHDRBS1-ATAAAA[ACG]
-log(pVal) up vs. bg
-log(pVal) dn vs. bg
Upregulated(519)
Downregulated(346)
Background(11780)
Motif Score (mean)
Negative log10(pValue)
0.035
0.026
0.018
0.009
0.0
4.054
3.040
2.027
1.013
0.0
0 50
-50 0 125 250
-250 -125 0 50
-50 0 50
-50 0

**Supplementary figure 6: Analysis of PTB motif enrichment after PTB KD. A:** Heatmap of the top enriched motifs in the Up SE events. Motif-map analysis of PTB motif enrichment in RI (**B**), MXE (**C**), A5SS (**D**), and A3SS (**E**) events.

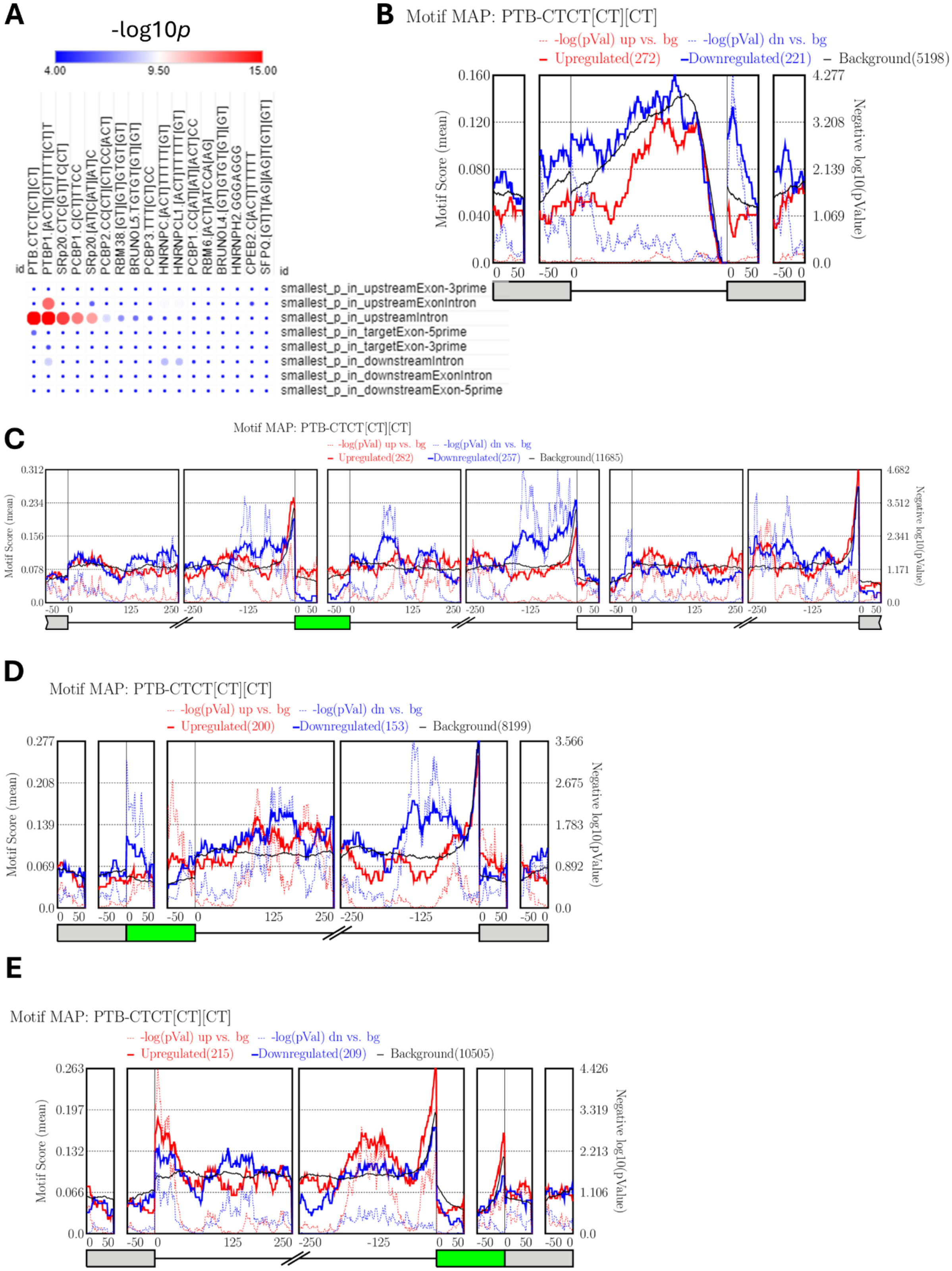
A
-log10p
4.00
9.50
15.00
id
PTB.CTCT[CT][CT]
PTBP1.[ACT][CT]TT[CT]T
SRp20.CTC[GT]TC[CT]
PCBP1.C[CT]TTCC
SRp20.[AT]C[AT][AT]C
PCBP2.CC[CT][CT]CC[ACT]
RBM38.[GT][GT]GTGT[GT]
BRUNOL5.TGTG[GT][GT]
PCBP3.TTT[CT]CC
HNRNPC.[ACT]TTTTT[GT]
HNRNPCL1.[ACT]TTTTT[GT]
PCBP1.CC[AT][AT][ACT]CC
RBM6.[ACT]ATCCA[AG]
BRUNOL4.[GT]GTGT[GT][GT]
HNRNPH2.GGGAGGG
CPEB2.C[ACT]TTTTT
SFPQ.[GT]T[AG][AG][GT][GT]
smallest_p_in_upstreamExon-3prime
smallest_p_in_upstreamExonIntron
smallest_p_in_upstreamIntron
smallest_p_in_targetExon-5prime
smallest_p_in_targetExon-3prime
smallest_p_in_downstreamIntron
smallest_p_in_downstreamExonIntron
smallest_p_in_downstreamExon-5prime
B
Motif MAP: PTB-CTCT[CT][CT]
-log(pVal) up vs. bg
-log(pVal) dn vs. bg
Upregulated(272)
Downregulated(221)
Background(5198)
Motif Score (mean)
Negative log10(pValue)
C
Motif MAP: PTB-CTCT[CT][CT]
-log(pVal) up vs. bg
-log(pVal) dn vs. bg
Upregulated(282)
Downregulated(257)
Background(11685)
Motif Score (mean)
Negative log10(pValue)
D
Motif MAP: PTB-CTCT[CT][CT]
-log(pVal) up vs. bg
-log(pVal) dn vs. bg
Upregulated(200)
Downregulated(153)
Background(8199)
Motif Score (mean)
Negative log10(pValue)
E
Motif MAP: PTB-CTCT[CT][CT]
-log(pVal) up vs. bg
-log(pVal) dn vs. bg
Upregulated(215)
Downregulated(209)
Background(10505)
Motif Score (mean)
Negative log10(pValue)

**Supplementary figure 7: A:** Venn diagram showing the overlap between upregulated SE events/genes after PTB KD and after neuronal differentiation. **B:** ORA analysis of the shared genes between PTB KD and neuronal differentiation from **A.**

A

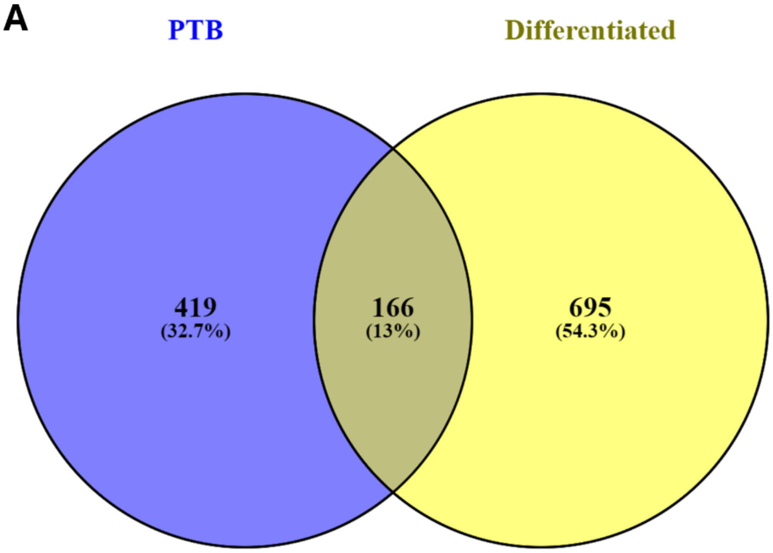
PTB
Differentiated
419
(32.7%)
166
(13%)
695
(54.3%)

B

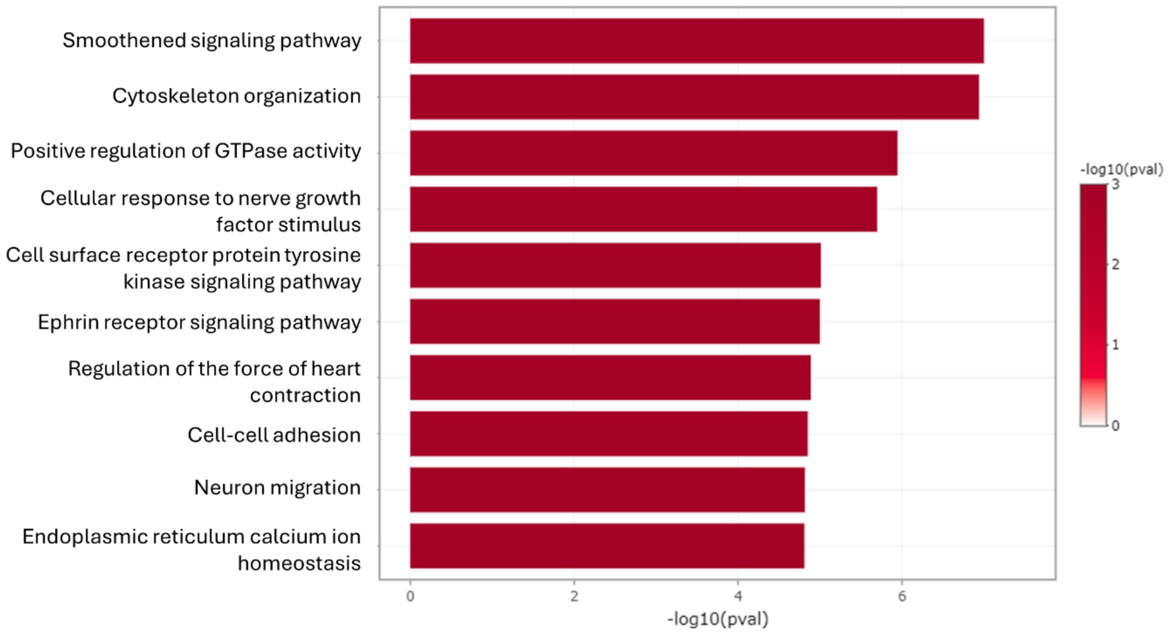
Smoothened signaling pathway
Cytoskeleton organization
Positive regulation of GTPase activity
Cellular response to nerve growth factor stimulus
Cell surface receptor protein tyrosine kinase signaling pathway
Ephrin receptor signaling pathway
Regulation of the force of heart contraction
Cell-cell adhesion
Neuron migration
Endoplasmic reticulum calcium ion homeostasis
0
2
4
6
-log10(pval)
-log10(pval)
3
2
1
0

**Supplementary figure 8:** Motif-map analysis of HuR motifs in SE events after HuR KD (continuation of figure 8C)

Motif MAP: HuR-TT[AGT]TTTT

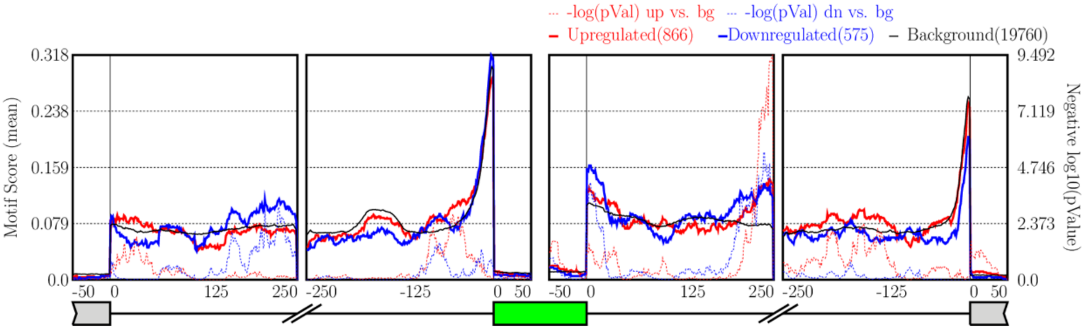


Motif MAP: HuR-TTT[AG][GT]TT

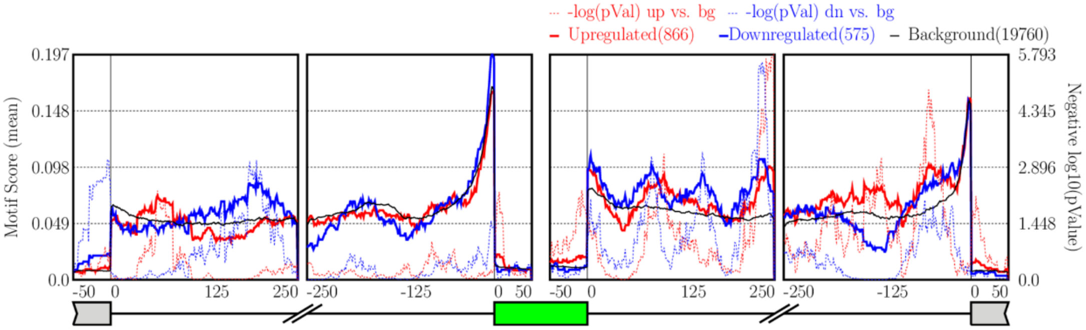


Motif MAP: HuR-TTTTTT[GT]

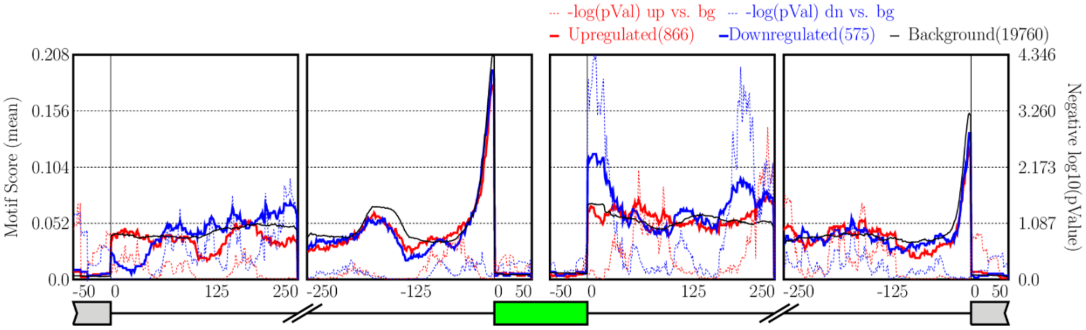


Motif MAP: HuR-TT[GT][AG]TTT

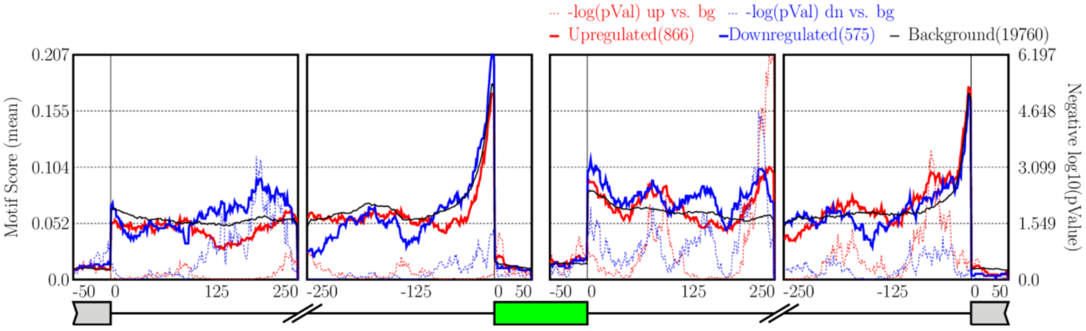

**Supplementary figure 9:** Motif-map analysis of HuR motifs in MXE events after HuR KD.

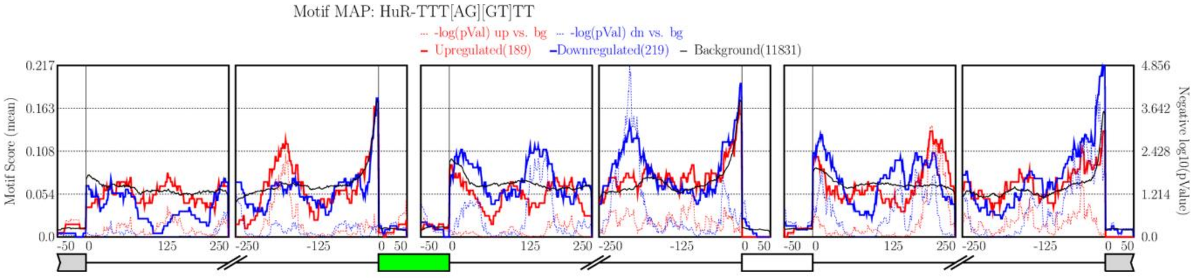
Motif MAP: HuR-TTT[AG][GT]TT
-log(pVal) up vs. bg
-log(pVal) dn vs. bg
Upregulated(189)
Downregulated(219)
Background(11831)
Motif Score (mean)
Negative log10(pValue)
0.217
0.163
0.108
0.054
0.0
4.856
3.642
2.428
1.214
0.0
-50 0 125 250 -250 -125 0 50

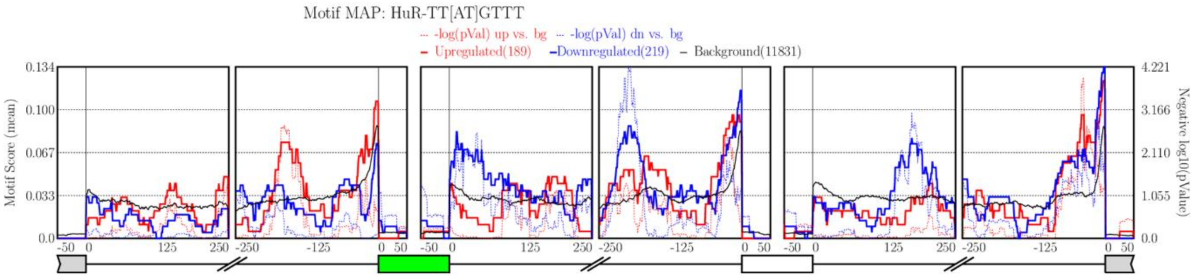
Motif MAP: HuR-TT[AT]GTTT
-log(pVal) up vs. bg
-log(pVal) dn vs. bg
Upregulated(189)
Downregulated(219)
Background(11831)
Motif Score (mean)
Negative log10(pValue)
0.134
0.100
0.067
0.033
0.0
4.221
3.166
2.110
1.055
0.0
-50 0 125 250 -250 -125 0 50

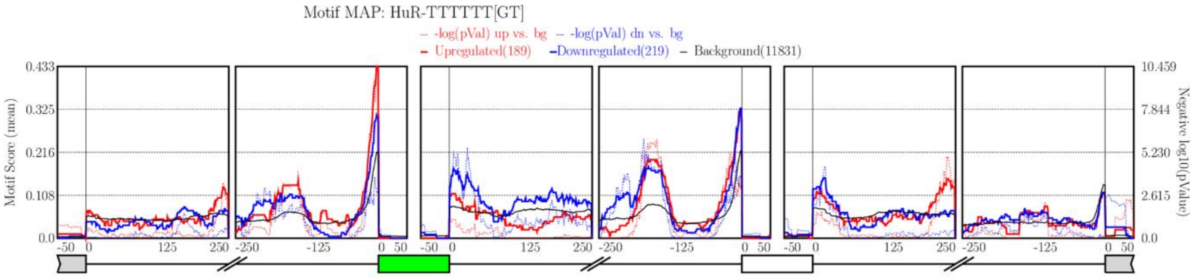
Motif MAP: HuR-TTTTTT[GT]
-log(pVal) up vs. bg
-log(pVal) dn vs. bg
Upregulated(189)
Downregulated(219)
Background(11831)
Motif Score (mean)
Negative log10(pValue)
0.433
0.325
0.216
0.108
0.0
10.459
7.844
5.230
2.615
0.0
-50 0 125 250 -250 -125 0 50

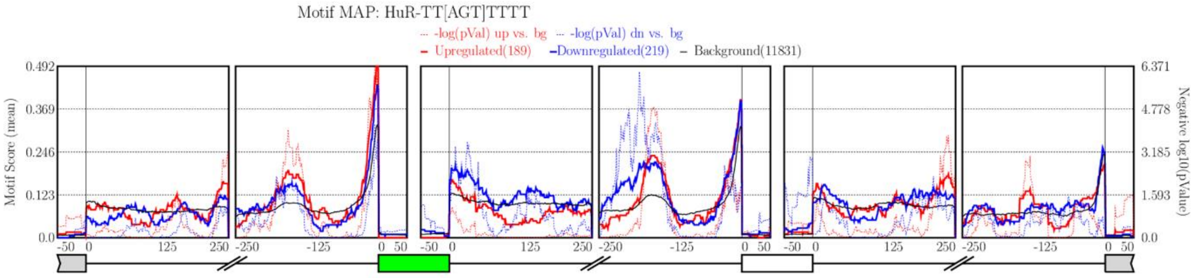
Motif MAP: HuR-TT[AGT]TTTT
-log(pVal) up vs. bg
-log(pVal) dn vs. bg
Upregulated(189)
Downregulated(219)
Background(11831)
Motif Score (mean)
Negative log10(pValue)
0.492
0.369
0.246
0.123
0.0
6.371
4.778
3.185
1.593
0.0
-50 0 125 250 -250 -125 0 50

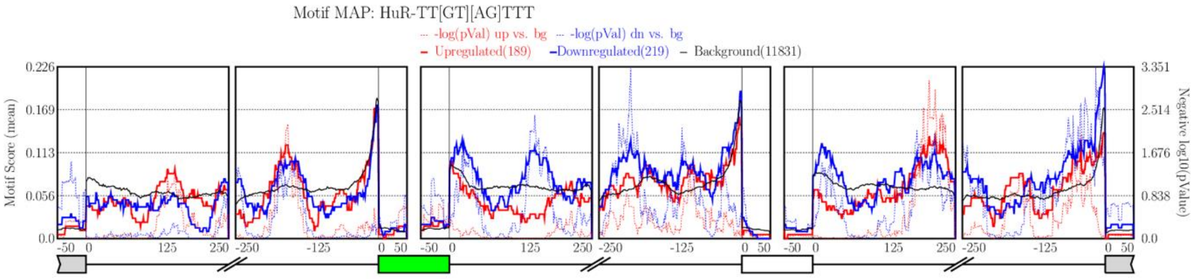
Motif MAP: HuR-TT[GT][AG]TTT
-log(pVal) up vs. bg
-log(pVal) dn vs. bg
Upregulated(189)
Downregulated(219)
Background(11831)
Motif Score (mean)
Negative log10(pValue)
0.226
0.169
0.113
0.056
0.0
3.351
2.514
1.676
0.838
0.0
-50 0 125 250 -250 -125 0 50

**Supplementary figure 10:** Motif-map analysis of HuR motifs in RI events after HuR KD.

Motif MAP: HuR-TTT[AG][GT]TT

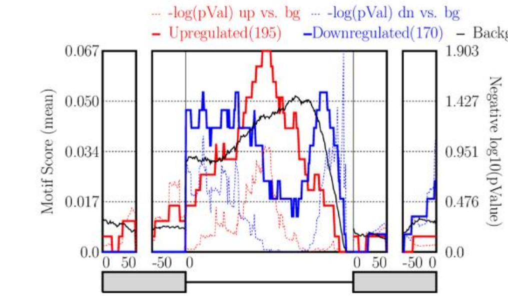


Motif MAP: HuR-TT[GT][AG]TTT

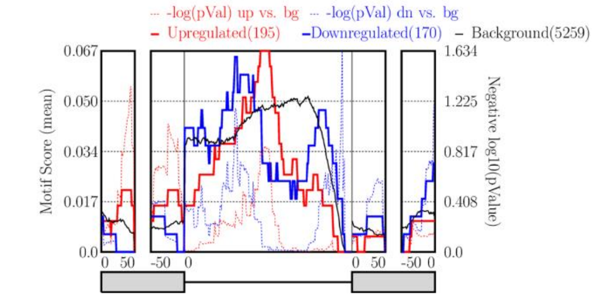


Motif MAP: HuR-TTTTTT[GT]

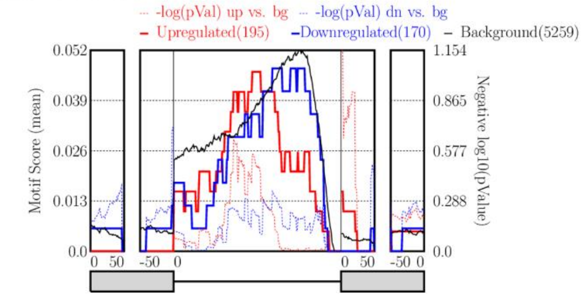


Motif MAP: HuR-TT[AGT]TTTT

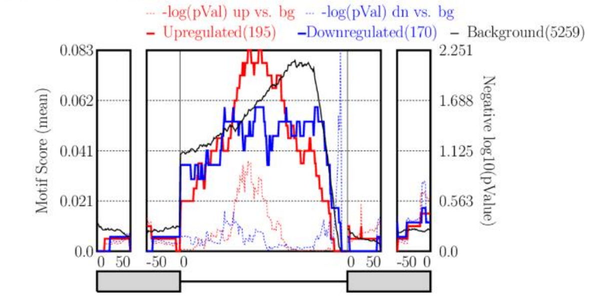


Motif MAP: HuR-TT[AT]GTTT

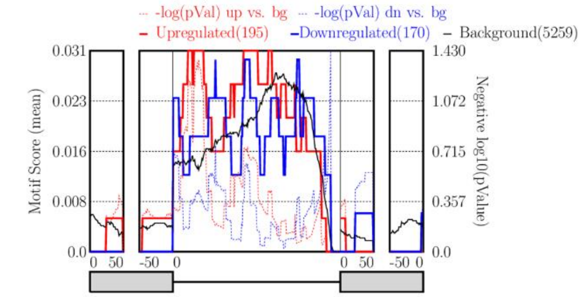

**Supplementary figure 11:** Motif-map analysis of HuR motifs in A5SS events after HuR KD.

Motif MAP: HuR-TTTTTT[GT]

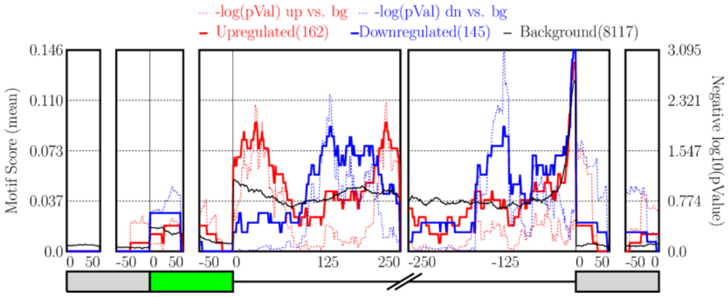


Motif MAP: HuR-TTT[AG][GT]TT

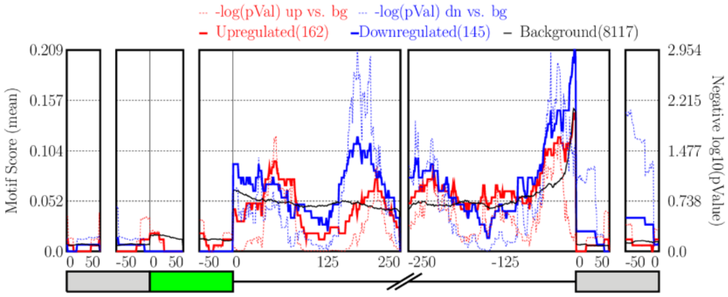


Motif MAP: HuR-TT[GT][AG]TTT

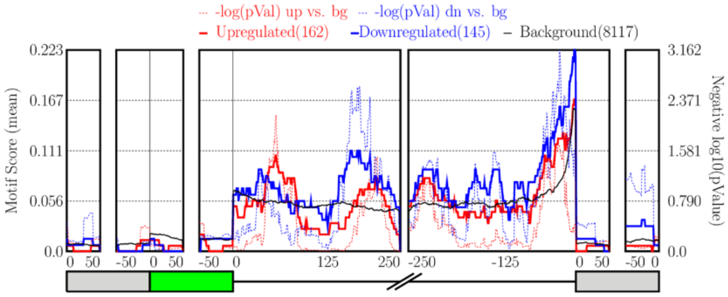


Motif MAP: HuR-TT[AGT]TTTT

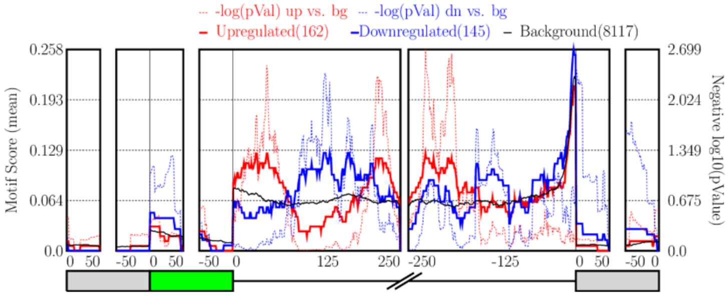


Motif MAP: HuR-TT[AT]GTTT

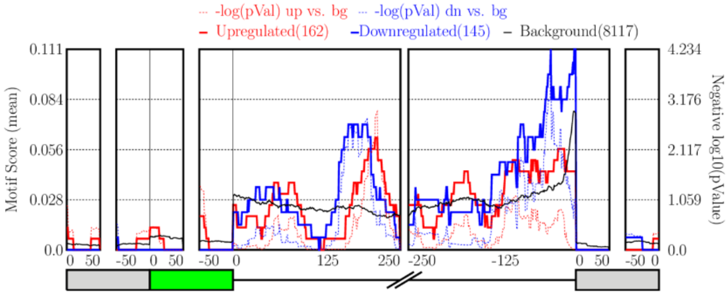

**Supplementary figure 12:** Motif-map analysis of HuR motifs in A3SS events after HuR KD.

Motif MAP: HuR-TTTTTT[GT]

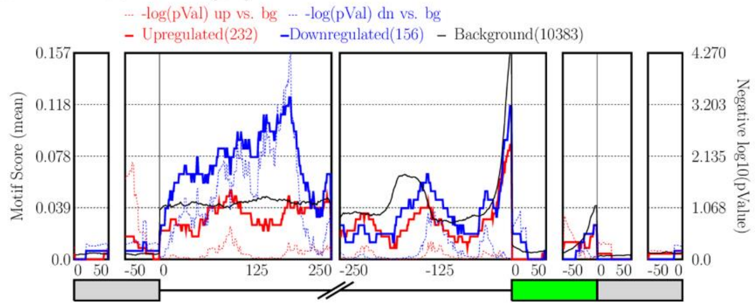


Motif MAP: HuR-TTT[AG][GT]TT

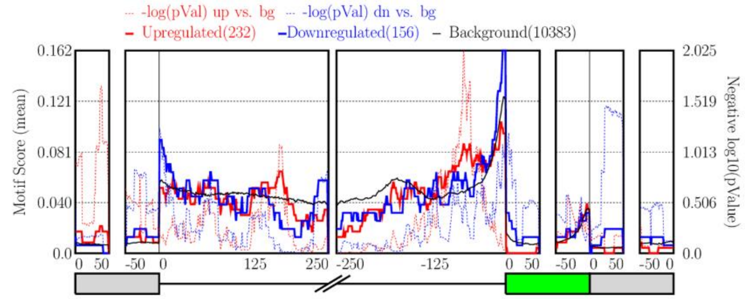


Motif MAP: HuR-TT[AT]GTTT

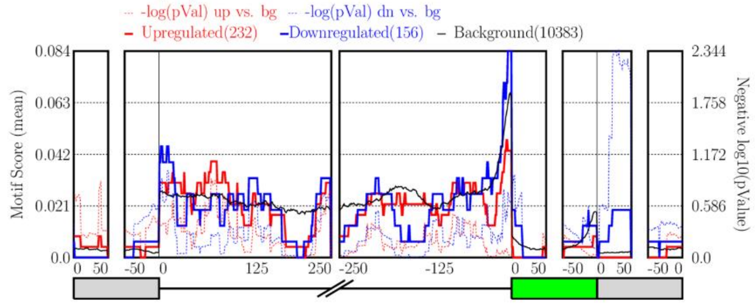


Motif MAP: HuR-TT[AGT]TTTT

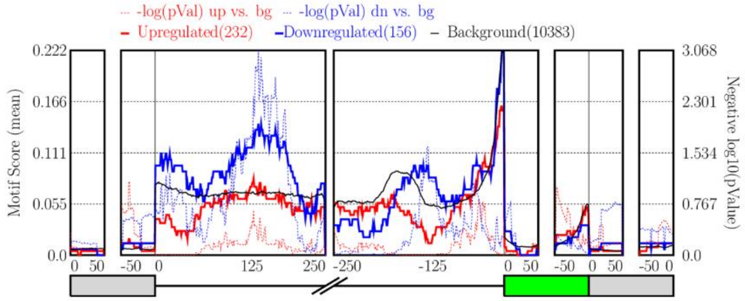


Motif MAP: HuR-TT[GT][AG]TTT

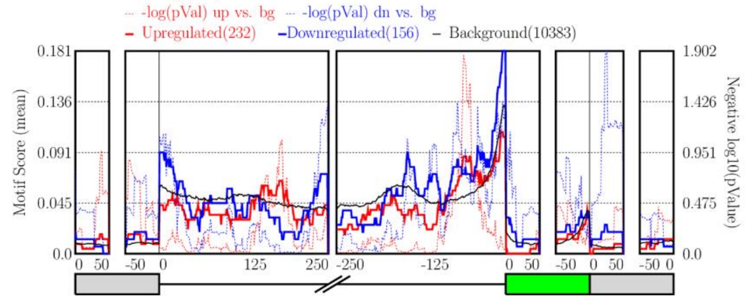

**Supplementary figure 13: A:** Venn diagram showing the overlap between downregulated SE events/genes after HuR KD and after neuronal differentiation. **B:** ORA analysis of the shared genes between HuR KD and neuronal differentiation from **A.**

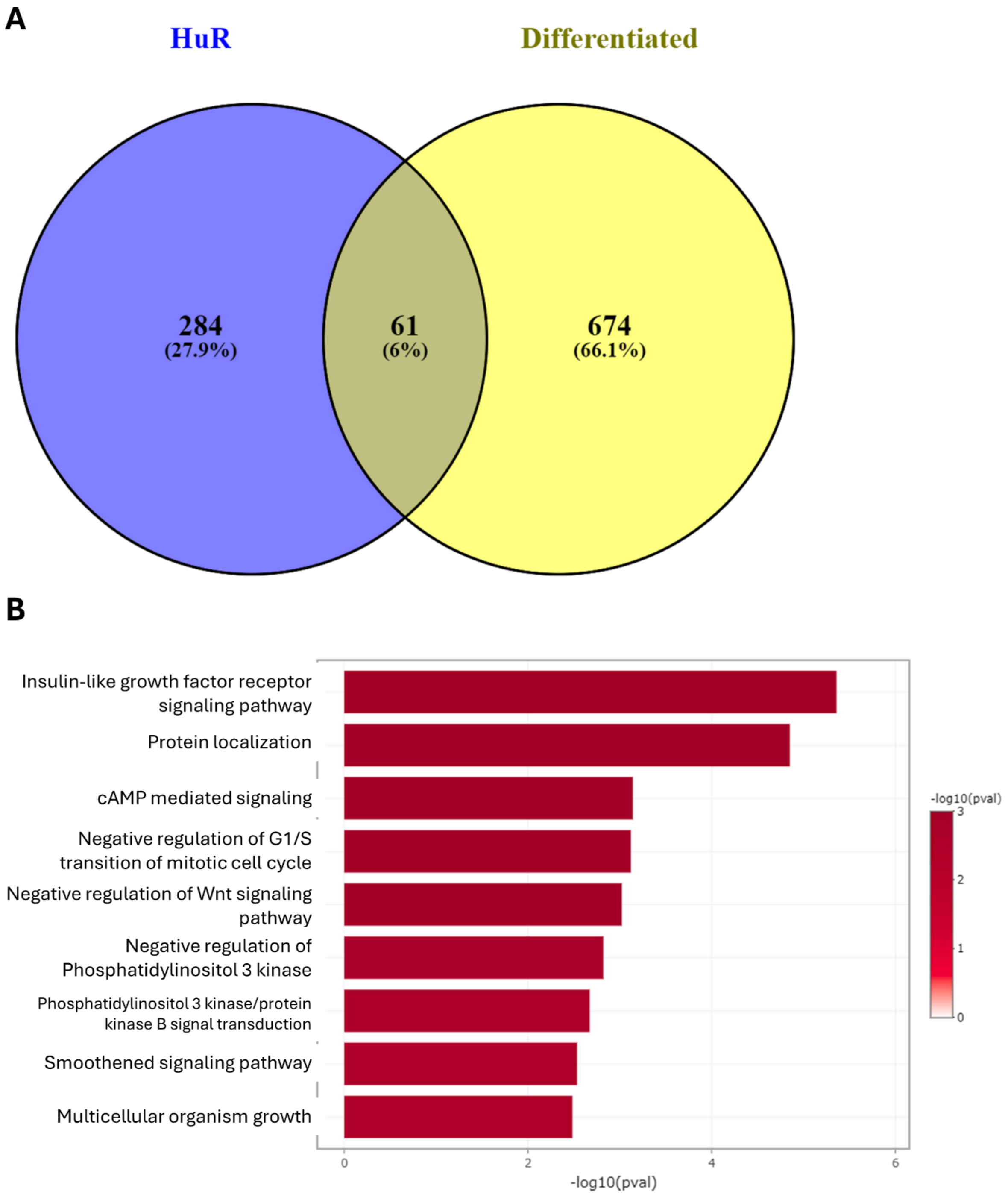
A
HuR
Differentiated
284
(27.9%)
61
(6%)
674
(66.1%)
B
Insulin-like growth factor receptor signaling pathway
Protein localization
cAMP mediated signaling
Negative regulation of G1/S transition of mitotic cell cycle
Negative regulation of Wnt signaling pathway
Negative regulation of Phosphatidylinositol 3 kinase
Phosphatidylinositol 3 kinase/protein kinase B signal transduction
Smoothened signaling pathway
Multicellular organism growth
-log10(pval)
0
2
4
6
-log10(pval)
3
2
1
0
